# Isotropic Reconstruction of Electron Tomograms with Deep Learning

**DOI:** 10.1101/2021.07.17.452128

**Authors:** Yun-Tao Liu, Heng Zhang, Hui Wang, Chang-Lu Tao, Guo-Qiang Bi, Z. Hong Zhou

**Author notes:** These authors contributed equally: Yun-Tao Liu, Heng Zhang. Correspondence should be addressed to: Z.H.Z, Or G.-Q. B.

## Abstract

Cryogenic electron tomography (cryoET) allows visualization of cellular structures *in situ*. However, anisotropic resolution arising from the intrinsic “missing-wedge” problem has presented major challenges in visualization and interpretation of tomograms. Here, we have developed IsoNet, a deep learning-based software package that iteratively reconstructs the missing-wedge information and increases signal-to-noise ratio, using the knowledge learned from raw tomograms. Without the need for sub-tomogram averaging, Isonet generates tomograms with significantly reduced resolution anisotropy. Applications of IsoNet to three representative types of cryoET data demonstrate greatly improved structural interpretability: resolving lattice defects in immature HIV particles, establishing architecture of the paraflagellar rod in Eukaryotic flagella, and identifying heptagon-containing clathrin cages inside a neuronal synapse of cultured cells. Therefore, by overcoming two fundamental limitations of cryoET, IsoNet enables functional interpretation of cellular tomograms without sub-tomogram averaging. Its application to high-resolution cellular tomograms should also help identify differently oriented complexes of the same kind for near-atomic resolution sub-tomogram averaging.

## Introduction

The advent of single-particle cryoEM has made it routine to determine structures of isolated macromolecular complexes at 2-4 Å resolution by averaging hundreds of thousands of particles, enabling atomic modeling. The biological functions of these complexes, however, are carried out through their interactions and often depend on their spatial arrangements within cells or sub-cellular organelles^1,2^. Examples abound, ranging from pleomorphic viruses, to cellular organelles, to large-scale cellular structures like synapses between neurons. Many viruses, notably those involved in devastating pandemics such as SARS-CoV-2, influenza viruses, and human immunodeficiency viruses (HIV), are pleomorphic in the organizations of their proteins and genomes. Cellular organelles, such as axonemes containing microtubule doublets surrounding a central pair^3^, though largely conserved in their core elements across different species, rely on their non-conserved and variable attachment of peripheral components that define their characteristic species-specific functions^4^. In neurons, organizations of molecules, rather than molecules alone, inside the synapse might underlie synaptic plasticity that is generally regarded as the cellular basis of learning and memory^5,6^. Such organizational information, or “molecular sociology”, unfortunately is lost in single-particle cryoEM.

To reveal such molecular sociology across viruses or inside cells, cryogenic electron tomography (cryoET) has become the tool of choice. This technique requires collecting a series images of the sample at different tilt angles, called “tilt series”. Due to radiation damage, limited electron dosage must be further fractionated throughout the tilt series, resulting in low signal-to-noise ratio (SNR) for the cryo tomogram. Furthermore, as tilting increases the effective thickness of the sample, the tilt range for cryoET is usually restricted to about ±70°. The missing views at higher tilt angles result in anisotropic resolution of the reconstructed 3D tomograms, with the resolution along the Z-axis the lowest (Supplementary Fig. 1). In Fourier space, these missing views lead to devoid of information in two continuous, opposing wedge-shaped regions, commonly referred to as the “missing-wedge”, along the tilt axis. This missing-wedge causes severe artifacts in 3D reconstruction of cellular cryoET, manifesting as, *e.g.*, oval-shaped synaptic vesicles^7^ (Supplementary Fig. 1). Thus, together with the low SNR in the reconstructed tomograms, the presence of missing-wedge artifacts prohibits direct interpretation of the reconstructed densities in 3D, which is key to the promise of cryoET to resolve molecule organization *in situ*.

Previous attempts have been made to partially recover information in the missing-wedge^8–10^ with *a priori* assumptions (*e.g.*, density positivity and solvent flatness) to constraint the structural features in reconstructed tomograms. However, such assumptions have limited information content (or “entropy”) and may not always hold true, given the complexity of biological systems. Alternatively, dual-axis tomography relies on imaging the same sample with two perpendicular tilt axes, reducing the two missing-wedges to two missing pyramids; thus it has the potential to alleviate artifacts in resulting tomograms^11^. However, acquisition and alignment of dual-axis tilt series are substantially more complicated than that of single-axis tilt series and could waste the already limited electron dose used for tilt series aquisition^12^. Consequently, dual-axis tomography, while implemented in high-end instruments such as the Thermo-Fisher Titan Krios, has not been practically attractive. Indeed, to date, no structure with better than nanometer resolution was obtained from dual-axis tomography.

Deep neural networks are known to learn relationships of complex data that are non-linear or have high dimensionality. In the field of computer vision, convolutional neural network (CNN) has been applied to various tasks, such as object recognition, image segmentation, and classification, often achieving high performance. In cryoEM field, CNN-based neural networks are applied to particle picking tasks and outperform conventional methods such as the Laplacian of Gaussian approach^13^. CNN is also introduced to cryoEM reconstruction to analyze heterogeneity of protein complexes with remarkable performance^14^. However, whether CNN can also recover missing-wedge information in cryoET has not been explored.

Here, we have developed a CNN-based software system, called *IsoNet*, for isotropic reconstruction of electron tomogram. IsoNet trains deep CNN that iteratively restores meaningful contents to compensate missing-wedge, using the information learned from the original tomogram. The resolution at Z-axis reaches about 30Å resolution as measured by the gold-standard Fourier shell correlation (FSC) criterion. By applying IsoNet to processing tomograms representing viral, organelle, and cellular samples, we demonstrate its superior performance in resolving novel structures of lattice defects in immature human immune-deficiency virus capsid, the scissors-stack-network architecture of the paraflagellar rod, and heptagon containing clathrin cage inside a neuronal synapse. The resulting tomograms with isotropic resolution from IsoNet should help direct interpretation and segmentation of 3D structure in cells and 3D picking hundreds of thousands of sub-tomogram particles for future near-atomic resolution cryoET studies.

## Results

### Workflow of IsoNet

In spite of anisotropic resolution, tomograms generated by cryoET reconstruction contain rich information with structural features such as plasma membranes, organelles, and protein complexes. Thus, it is possible to recover the missing information by merging information from similar features present in the same tomograms but at different orientations relative to each other. An example of filling such missing information is through subtomogram averaging, which aligns and averages structures of particles that are identified to be identical but at different orientations in the tomogram. IsoNet is designed to expand this technique to reconstruct missing-wedge information by training the neural network targeting the subtomograms at different rotations for both regular and polymorphous structures.

The workflow of IsoNet contains five steps (Fig. 1a). Among them, three are major and required: *Extract*, *Refine* and *Predict*; and the other two are optional: *Deconvolve CTF* and *Generate Mask*. Each of these steps can be performed with one command of IsoNet in Linux terminal. Among the 5 steps, *Refine* and *Predict* relies on graphical processing unit (GPU) that provides superior processing power. The input of IsoNet is either from a single or multiple tomograms. Based on the principle of machine learning, more tomograms will generate more reliable results but takes longer processing time. In practice, the typical number of tomograms for IsoNet is from one to five. The tomogram(s) can be reconstructed by either weighted back projection (WBP) or iterative methods, such as simultaneous iterative reconstructive technique (SIRT).

**Fig. 1.**
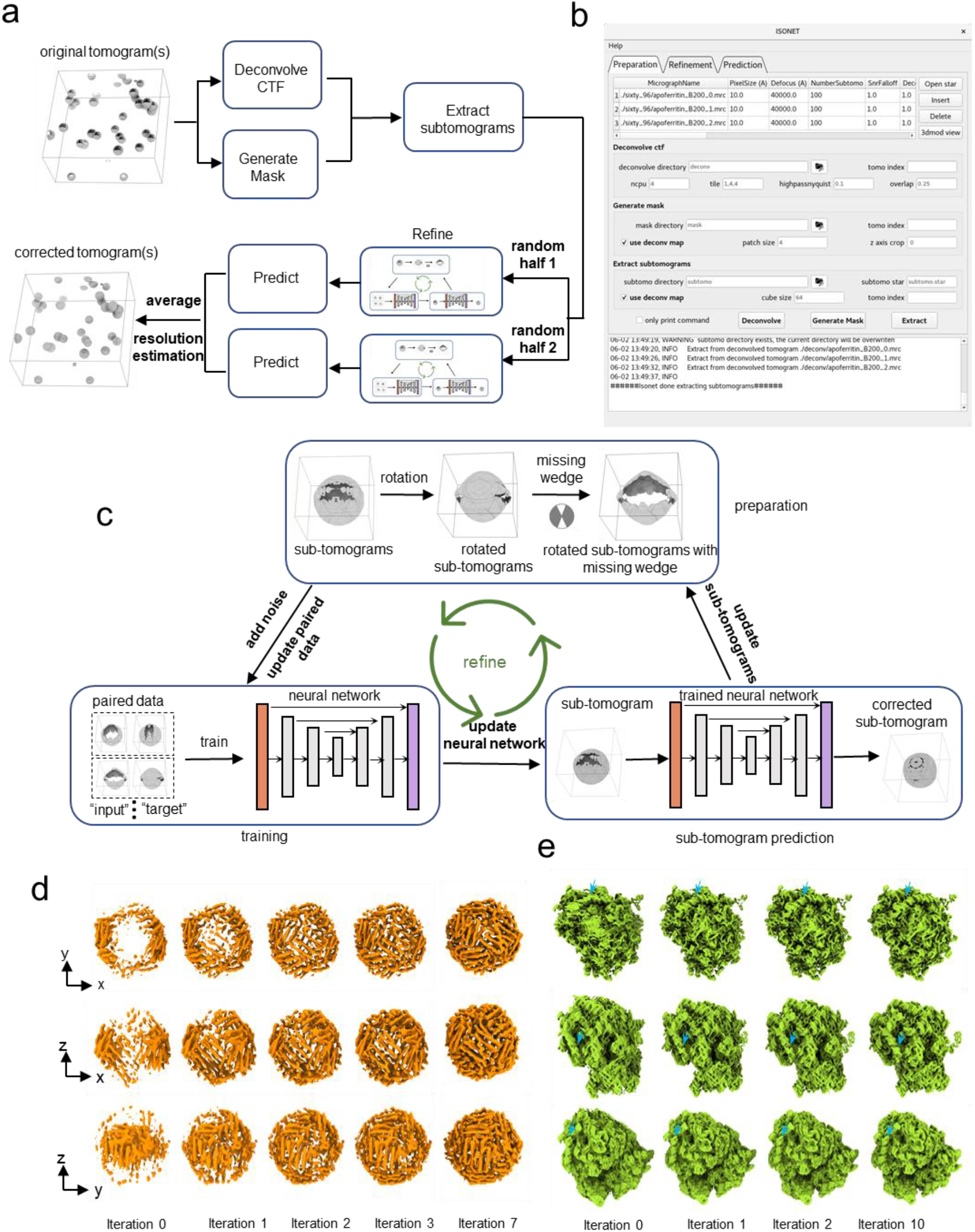
Principle and workflow of IsoNet. **a,** Workflow of the IsoNet software. **b,** GUI of IsoNet. **c,** Illustration of *Refine* step: First, subtomograms are rotated and then applied with additional missing-wedge artifacts to produce paired data for training. Second, the paired data is used to train a neural network with U-net architecture. Third, the trained neural network is applied to the extracted subtomograms to produce missing-wedge corrected subtomograms. The recovered information in these subtomograms is added to the original subtomograms, producing new datasets for the next iteration. **d**, Validation of IsoNet with simulated sub-tomogram of apoferritin and ribosome. Surface views from three orthogonal directions of the reconstructions are shown after increasing iterations of IsoNet processing. Blue arrows indicate segments of RNA duplexes.

We implemented IsoNet in Python using Linux as the native operating system. The package takes advantage of the Keras interface of well-established Tensorflow platform^15^ and can be run standard-alone, independent of other software packages. The package can be run either through command line or through a graphical user interface (GUI) (Fig. 1b), thus meeting to the needs of both seasoned and novice cryoET investigators. The GUI contains three tabs to facilitate navigation. In each tab, information of the tomograms and the parameters for each command can be specified. By clicking “*Deconvolve*”, “*Generate Mask*”, “*Extract*”, “*Refine*” and “*Predict*” buttons, user can execute the corresponding command. The “only print command” option prints out the corresponding command for each step which can be executed on other computers or submitted to computer clusters.

#### Deconvolve CTF and Generate Mask steps

These two optional steps are performed on the input tomograms prior to the subtomogram extraction in *Extract* step (Fig. 1a). The *Deconvolve CTF* step has two purposes: to enhance low-resolution information and compensate for the contrast transfer function (CTF) in the tomograms acquired at certain underfocus conditions. Due to the presence of zeros in CTF, we used a Weiner filter for CTF compensation, as implemented in Warp^16^. The *Generate Mask* step uses statistical methods to detect “empty” areas in the tomograms (including vacuum above and below the sample and those only containing ice or carbon) to be excluded from the subsequent analysis. Both steps could improve performance and efficiency of neural network training.

#### Extract step

This step allows randomly cropping subtomograms in the original tomograms or the region-of-interest of the tomograms defined by masks. The maximum sizes of subtomograms depend on the memory of graphics processing units (GPU), and 64^3^ or 96^3^ voxels are often used. The extracted subtomograms can be split into random halves to train the neural network independently (Fig. 1a), allowing users to perform 3D gold-standard FSC^17,18^ to determine the resolution of IsoNet reconstructed tomograms over different angular directions, particularly on Z-axis.

#### Refine Step

Central to the IsoNet workflow is the *Refine* step, which iteratively trains neural networks to perform missing-wedge correction and denoising (Fig. 1c). Training of the neural network requires paired subtomograms as the “inputs” and the “targets.” The “targets” for IsoNet are the extracted subtomograms rotated at different orientations. In total, 20 different orientations are defined in IsoNet, generating 20 “target” subtomograms for each extracted subtomogram (Supplementary Fig. 2). For each “target” subtomogram, the missing-wedge is computationally imposed in Fourier space to generate the corresponding “input” subtomograms (Fig. 1c). After generating the paired dataset, we train a neural network to map the “input” to the “target”, enabling the network to recover the imposed missing-wedge artifacts. The neural network used in IsoNet adopts U-Net architecture^19^, containing an encoder path that extracts low-dimensional representation retaining essential properties, a decoder path to reconstruct from the encoded representation, and skip-connections between encoder and decoder to preserve high-resolution information (Supplementary Fig. 3).

However, the “target” in the data pairs described above are not ideal subtomograms. These subtomograms, though rotated, still miss information in other directions. To recover that information and make “target” subtomograms resembling “ground truth”, we adopt an iterative approach: In the first iteration, we train the network with subtomograms generated from the *Extract* step and obtain the IsoNet-predicted subtomograms. Then, the gained information in the missing-wedge region in the Fourier space of the predicted subtomograms was added to the original subtomograms, generating the first-iteration missing-wedge corrected subtomograms (Fig. 1c). To further improve miss-wedge correction with more iterations, the corrected subtomograms from the previous iteration are used for the paired data generation in the next iteration because they are closer to missing-wedge-free 3D volumes than the extracted original subtomograms. The trained network from the previous iteration is then refined with the newly generated data pairs. Through multiple iterations, the missing-wedge information is gradually added to the subtomograms (Fig. 1c and Supplementary Fig. 4). Usually, after 10-20 iterations, the refinement converges when the mean square error no longer decreases. The result of this *Refine* step is a trained network that will be applied to the full tomograms and produce the isotropic reconstruction in the *Predict* step (Fig. 1a).

Within the *Refine* step of IsoNet, we also implemented a denoising module based on the noisier-input strategy^20,21^. When this optional denoising module is enabled in the *Refine* step, in each iteration, 3D noise volumes are reconstructed by the back-projection algorithm from a series of 2D images containing only Gaussian noise. Those 3D noise volumes are then added to “input” subtomograms, with the “target” subtomograms staying the same. With this strategy, IsoNet can be robustly trained with these noisier “input” subtomograms to eliminate the added noise and improve the SNR of final isotropic reconstructions (Fig. 1c and Supplementary Fig. 4).

#### Predict step

This step performs missing-wedge correction by applying networks obtained in *Refine* step to the tomograms of interest. This *Predict* step runs much faster than the *Refine* step. The tomograms used for *Predict* step are typically (preferably because there are no concerns of bias) the same or a subset of the tomograms used to train the network. Nonetheless, users can in theory apply the trained network to tomograms of other similar samples.

### Benchmarking with simulated data

We first perform IsoNet reconstruction on simulated subtomograms using the public available atomic models. Two scenarios have been considered: apoferritin^22^ for the first test because it has been widely used as a benchmarking specimen in high-resolution cryoEM and ribosome^23^ as the second test due to its asymmetric shape and primarily nucleic acid content. For both tests, density maps are simulated from the atomic models using *molmap* function in Chimera^24^ and filtered to 8Å resolution (Figs. 1d and e). The simulated maps were then rotated in 10 random directions and imposed missing-wedge in Fourier space, resulting in simulated subtomograms with missing-wedge artifacts (leftmost columns in both Figs. 1d and e).

As evident in both tests with simulated subtomograms, features such as alpha-helices perpendicular to the Z-direction are smeared out in those simulated subtomograms due to the missing-wedge artifact. IsoNet was then used to process those simulated subtomograms. As expected, the missing information was recovered during this iterative refinement process (Figs. 1d and e). After 7 iterations, all the alpha helices are visible and identical to the ground truth structures in the first test. The cubic symmetry of apoferritin gradually emerged even though we did not impose symmetry during the processing using IsoNet. In the second test, the distortion in the shapes of ribosome is reduced during the *Refine* step, with the major and minor grooves of the RNA become distinguishable (Fig. 1e). These results indicate that IsoNet performed well with simulated round/symmetric protein complex as well as asymmetric complex containing both protein and nucleic acid.

### Application to virus tomograms

To further demonstrate the superior performance of IsoNet in real-world examples, we perform the IsoNet reconstruction with the well-characterized cryoET datasets of virus-like particles (VLP) of immature HIV-1, which is publicly available from the Electron Microscopy Pilot Image Archive^25,26^ (EMPIAR-10164).

After reconstructed with IsoNet, gold beads in the tomogram appear spherical (Fig. 2a), as they should, instead of the “X” shape due to the missing-wedge problem. Notably, the top and the bottom of the VLP can now be observed in the IsoNet generated tomogram. When examined in the Fourier space, the missing-wedge region on the XZ slices was filled with values compared to the Fourier transform of the original tomogram devoid of the information (Fig. 2a). To quantify the resolution of the filled information, we spilt the extract subtomograms into two random subsets, trained two neural networks using those two subsets independently, and then performed 3D FSC calculation^17^. The resolution on the XY plane is higher than other planes (Fig. 2b), with the resolution along the X and Y axis reaching the Nyquist resolution, showing our network preserves the high-resolution information of the original tomograms. The Z-axis resolution of the isotropic resolution is about 30Å (Fig. 2b), which is the lowest resolution in all directions. This result demonstrates that our isotropic reconstruction can faithfully reconstruct the missing-wedge information at least 30Å resolution.

**Fig. 2.**
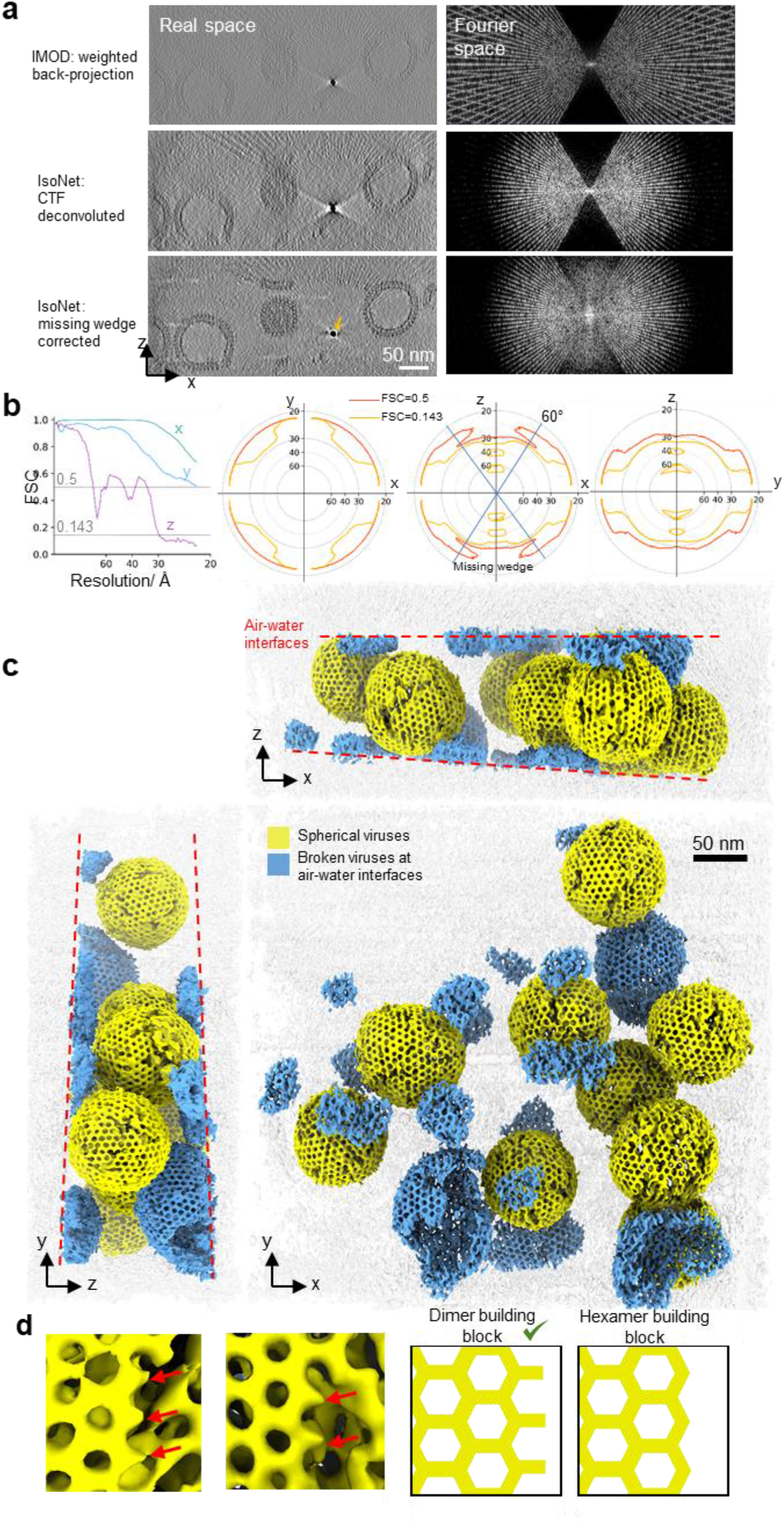
IsoNet reveals lattice defects in immature HIV capsid. **a,** XZ slice views of tomogram reconstructed with WBP (top), CTF deconvoluted WBP tomogram (middle), and missing-wedge corrected tomograms (bottom), with their Fourier transforms on the right. Orange arrow indicates a gold bead. **b,** 3D FSC of the two independent isotropic reconstructions, the left panel shows the FSC along the X, Y and Z directions. Three panels on the right show the 3D FSC at 0.5 and 0.143 cutoffs on XY, XZ, and YZ planes, respectively. **c,** 3D rendering of the missing-wedge corrected tomogram. Dashed lines show the air-water interfaces. **d,** Examples (left) and illustrations of the lattice edges of HIV capsids. Red arrows point out the density protrusions on the edges of hexagonal lattices.

Importantly, our isotropic 3D reconstruction shows that the quality of the structure is similar across all directions, allowing biological structures to be interpreted adequately (Fig. 2c and Supplementary Video 1). We resolved those broken viruses, sheared along top and bottom planes of the tomograms (Fig. 2c and Supplementary Video 2), indicating that the air-water interfaces caused deformation of the capsid, as well-recognized in the cryoEM field^27^. The denatured Gag proteins, which are subunits of capsids, at the air-water interfaces are mostly featureless.

The spherical viruses that were fully embedded in ice are made of hexagonal lattices (Fig. 2c), whereas no pentagon subunit is observed, consistent with the subtomogram averaging results of immature HIV particles^26^. Lattice defects are incorporated onto the hexagonal lattices, making gaps between patches of the lattices (Fig. 2c). These defects and slight curvature on the hexagonal lattices could enable the formation of the spherical shape without pentagons. On lattice edges, small density protrusions extending from the hexagons were observed (Fig. 2d), indicating the complete hexagons are not assembly units of HIV. In concert with this observation, a recent study shows the Gag dimers are the basic assembly units of the HIV-1 particle^28^. These protrusions could be Gag dimers and are prone to structural changes during proteolytic cleavage^28^. Those 3D details on HIV lattices can only be directly visualized after processed by IsoNet. Taken together, the above observations demonstrate that IsoNet can effectively compensate for the missing-wedge problem for relatively thin but heterogeneous structures, such as the immature HIV particles, and reach about 30Å Z-axis resolution.

### Application to tomograms of cellular organelles

Next, we tested the performance of IsoNet on resolving structures within cellular organelles by processing tomograms of flagella of *Trypanosoma. Brucei* using IsoNet. The missing-wedge compensated tomogram shows relatively uniform or isotropic structures, in all three dimensions (Figs. 3a and b). The overall contrast is better than the original tomogram partially due to the denoising of the network. One noticeable missing-wedge artifact is that it is difficult to recognize the well-established 9 (outer doublets) + 2 (central-pair singlets) microtubule arrangement in the cross-section view (i.e., XZ view in Fig. 3a). This arrangement can be readily visible in the result generated by IsoNet (Fig. 3b). The missing-wedge effect is also reflected by the broken and oval-shaped microtubules and severe artifacts in XZ and YZ planes in the original tomogram reconstructed with SIRT algorithm (Fig. 3a). In tomograms generated by IsoNet, the microtubules become complete and circular-shaped with some visible tubulin subunits (Fig. 3b and Supplementary Fig. 5). Binding to the microtubules, the arrays of outer (red arrows in Fig. 3b) and inner (blue arrows in Fig. 3b) arm dynein proteins are now clearly distinguishable in the IsoNet generated tomogram. And radial spokes connecting the outer doublets to the central pair can be distinguished in all three orthogonal slices (Fig. 3a-b).

**Fig. 3.**
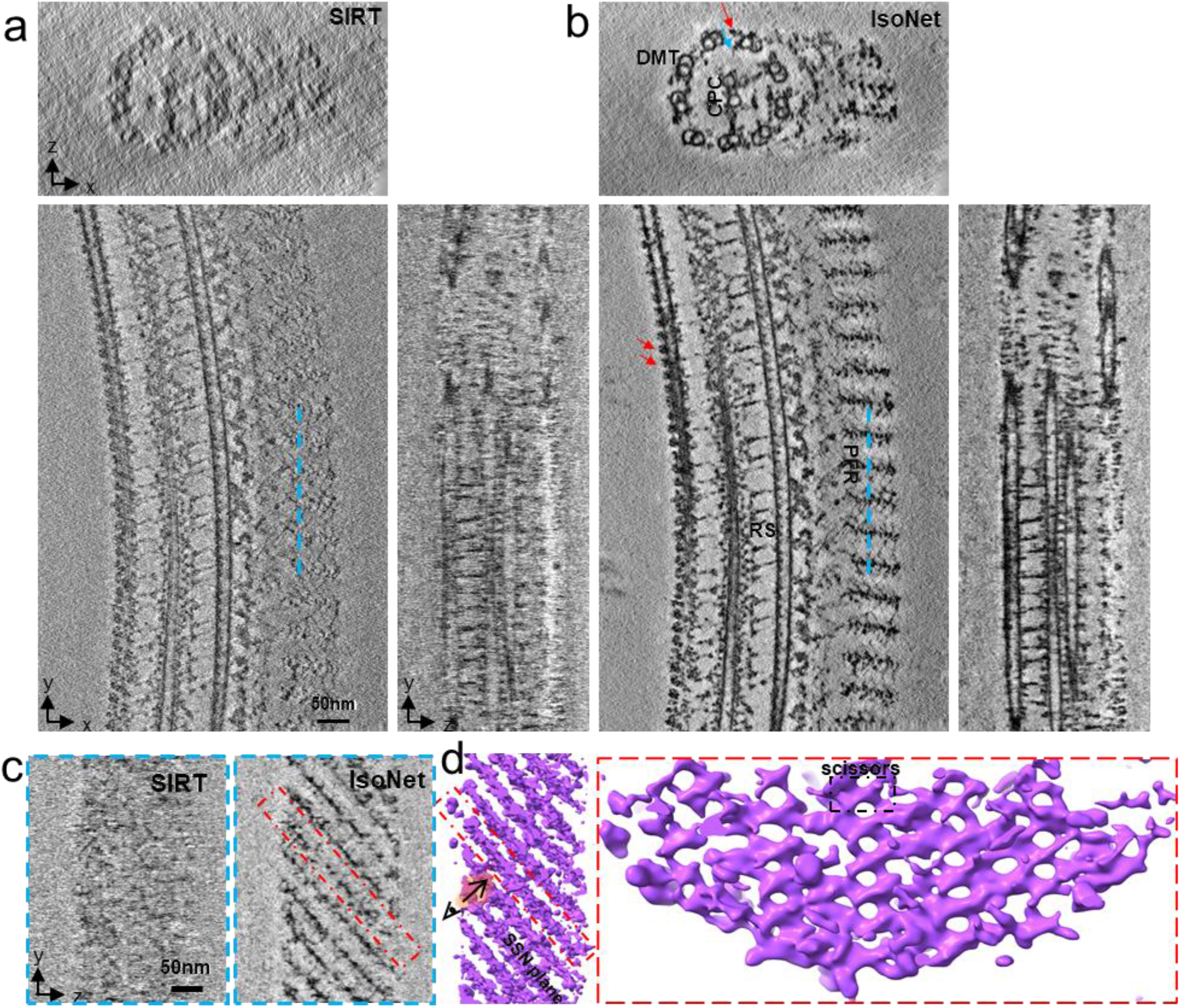
Architecture of the PFR revealed after missing-wedge correction. **a-b,** Orthogonal slices of a tomogram of flagellar for SIRT reconstruction **(a)** and IsoNet reconstruction **(b)**. DMT: Doublet microtubule; CPC: central pair complex; RS: radial spoke; Red arrows: outer arm dyneins; Blue arrow: inner arm dynein. **c,** YZ slices show the cross-sections corresponding to the cyan lines in **(a)** and **(b)**. **d,** 3D rendering of PFR in the IsoNet generated tomogram. Left panel is the 3D view of PFR in the direction corresponding to **(c)**. Right panel shows the *en face* view of a PFR scissors-stack-network (SSN) plane.

On one side of 9+2 microtubules lies paraflagellar rod (PFR). The structure of PFR is obscure in the tomogram reconstructed by SIRT (Fig. 3c), which has given rise to the long-lasting debate of the PFR organization^29–31^. The IsoNet generated tomograms showed a much clearer picture of PFR. PFR density consists of parallelly arranged planes, and the angle between those planes and the direction of the axoneme is 45°. Within these planes, scissors-like densities stack upon each other, forming a scissors-stack-network (Figs. 3d). This highly organized mesh structure could serve as a biological spring to assist the movement of the flagella. This unique PFR structure observed here is consistent with the organization resolved through tedious efforts of sub-tomogram averaging of thousands of sub-tomograms^32^. The above observations demonstrate that IsoNet can compensate for the missing-wedge problem for nonspherical cellular organelles, such as those in the Eukaryotic flagella, and unveil structure with meticulous details without the need of sub-tomogram averaging.

### Applications to tomograms of cells

To evaluate IsoNet’s performance for much larger and more complex structures in cells, we applied IsoNet to tomograms of synapses in cultured hippocampal neurons^7^. Hippocampal synapses are key devices in brain circuits for information processing and storage. They are about 200-1000 nm in size, rich in proteins, lipid membranes, vesicles, mitochondria, and other organelles^7,33,34^. These samples are thicker^7^ (300-500 nm) than the above-described flagella and virus samples, thus are representative of low SNR tomograms. The intrinsic molecular crowding and structural complexity of the synapse also present difficulties for missing-wedge correction. Arguably, synaptic cryoET tomograms are among the most challenging datasets for any analysis algorithm.

IsoNet achieved isotropic reconstruction of the synaptic tomogram with substantially higher contrast and better structural integrity (Figs. 4a, b, and Supplementary Videos 3-5). Synaptic vesicles that were smeared out along the Z-axis in the original tomograms now become spherical (Fig. 4c). The linker proteins between vesicles that are hardly seen in the original tomograms now become visible in XZ and YZ planes (Figs. 4c). Even some horizontally oriented features can be resolved. For example, large patches of membrane on the top and the bottom of the synapse and the endoplasmic reticulum (ER) now appear smooth and continuous in the isotropic reconstruction (Figs. 4b and e). These structural integrity improvements facilitate the segmentation of the cellular structure, since the missing-wedge corrected structures can be directly displayed based on their density threshold in 3D. Particularly, placing the artificial spheres to represent synaptic vesicles, as in previous studies^7,33^, is no longer needed (Fig. 4e). As the elongation effect of microtubules in the Z-axis being corrected, the protofilaments of microtubules have now become visible (Figs. 4d and f). Inside synapses, numerous small black dots can be observed in the cytoplasm but not in vesicular lumens. These dots represent small cytoplasmic proteins (orange arrows in Fig. 4c), indicating our reconstruction preserves delicate structural features.

**Fig. 4.**
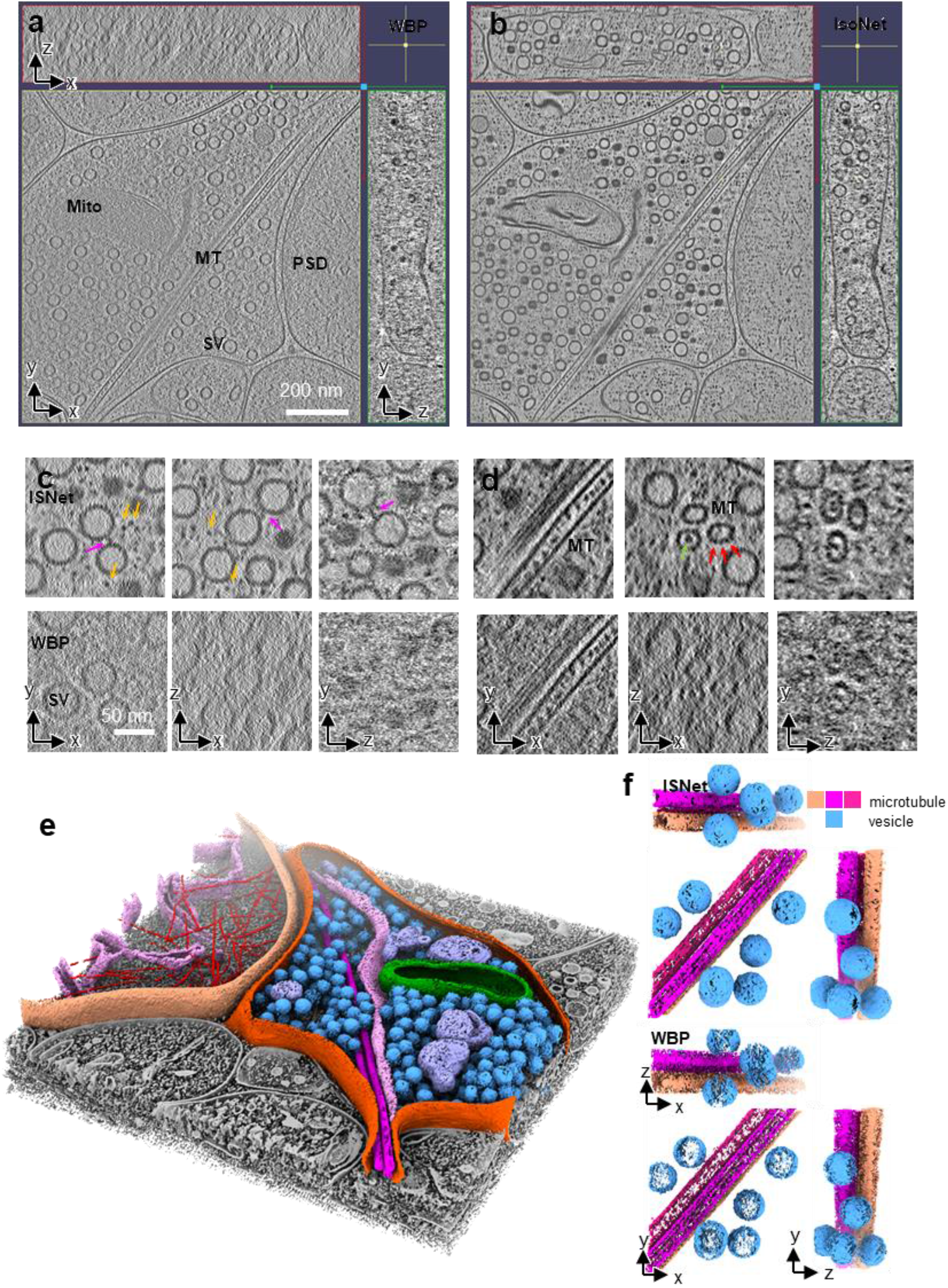
IsoNet recovers missing information in the tomograms of neuronal synapses. **a-b,** Orthogonal slices of a synaptic tomogram reconstructed with WBP **(a)** and IsoNet **(b)**. SV: synaptic vesicle; Mito: mitochondria; MT: microtubule; PSD: postsynaptic density. **c-d,** Zoomed-in orthogonal slices of WBP reconstruction and IsoNet produced reconstruction. Magenta arrows: vesicle linker; Orange arrows: small cellular proteins; Green arrow: microtubule luminal particles; Red arrows: microtubule subunits. **e,** 3D rendering of the tomogram shown in **(b)**. **f,** 3D rendering of a slab of tomogram with WBP reconstructions and Isotropic reconstructions, showing microtubules and vesicles.

As a prominent example, tomograms from IsoNet revealed various types of clathrin coats in hippocampal synapses. Clathrin-mediated endocytosis is a well-known presynaptic vesicle recycling mechanism and is a critical step in synaptic transmission^35,36^. Clathrin proteins are also present in postsynaptic compartment for neurotransmitter receptor endocytosis, a process playing essential roles in synaptic plasticity^37^. Those clathrin proteins are known to form cages that consist of pentagons and hexagons^38^. We observed structures similar to clathrins cages of various sizes in the postsynaptic compartment in synaptic cryo tomogram. However, due to the missing-wedge effect, the geometry of these clathrin cages cannot be directly resolved *in situ* in typical cellular tomograms. We applied IsoNet software to one synaptic tomogram that contains many putative clathrin cages in the postsynaptic compartment (Figs. 5a and b, Supplementary Fig. 6). After the isotropic reconstruction, all the pentagons and hexagons, which made up the clathrin cages, are revealed (Figs. 5c and d). This contrasts with the original tomograms, where the polygons are smeared, especially in XZ and YZ planes.

**Fig. 5.**
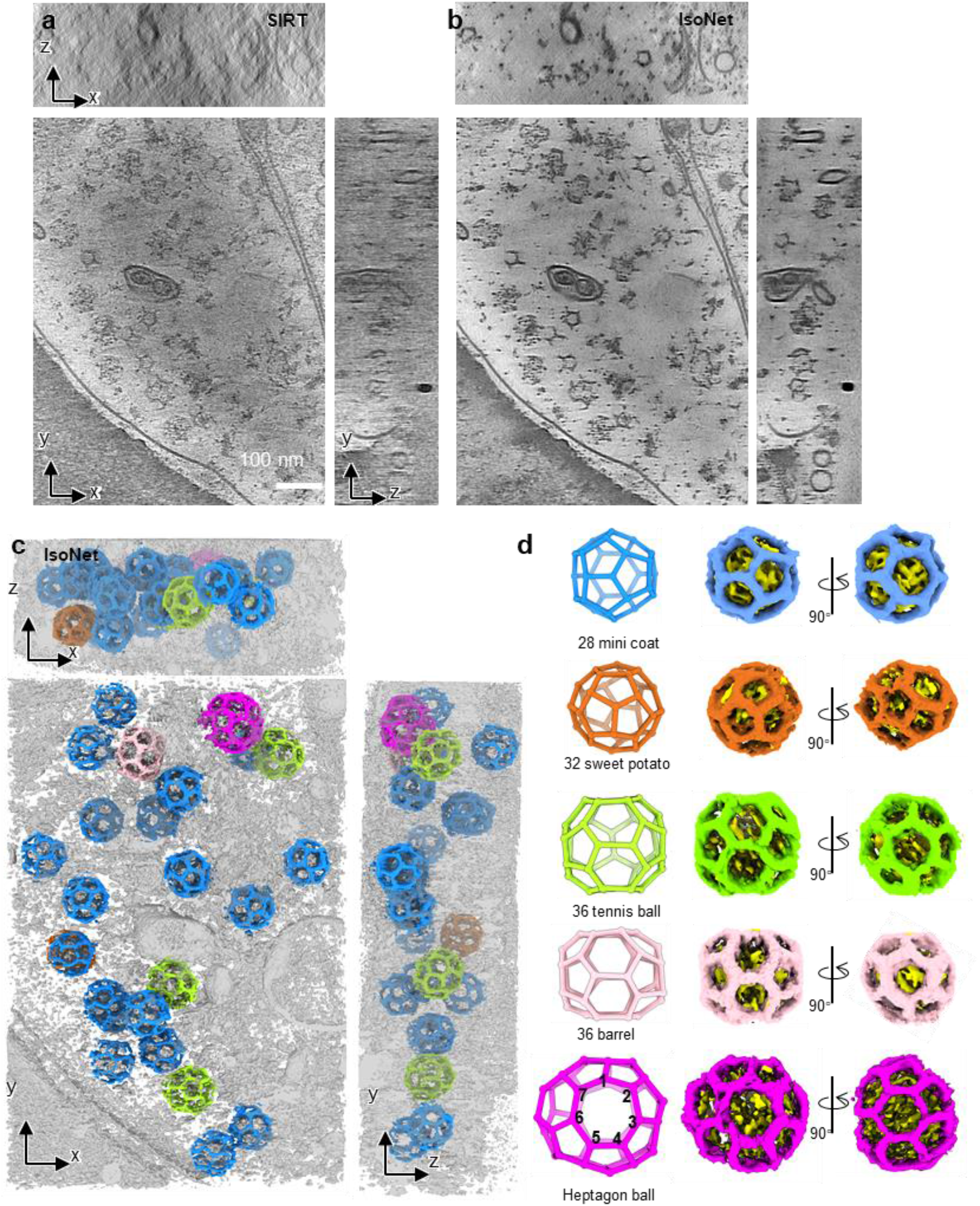
IsoNet reveals various types clathrin coats in a synapse. **a-b,** Orthogonal slice views of another synaptic tomogram reconstructed with SIRT algorithm **(a)** and IsoNet **(b)**. **c,** 3D rendering of the tomogram shown in (**b**). **d,** 3D views of the five types of clathrin cages in **(c)**.

The 25 clathrin cages can be categorized into five types based on their geometry (Supplementary Video 6). The most abundant type is minicoat, which is the smallest cages the clathrin proteins can form^39^. Intriguingly, the largest clathrin cage contains two heptagons, in addition to 8 hexagons and 14 pentagons (Fig. 5d, Supplementary Fig. 7), which has not been reported in previous single-particle analysis^38,39^. This geometry of the cage deviates from the common belief that a closed polyhedral protein cage contains 12 polygons. This heterogeneous in the Platonic cages of the clathrin arises from the specific yet variable forms of clathrin triskelion interactions. Adapting those heptagons in neurons could likely be strategy to scale up the size of the clathrin coats that enables accommodating different sizes of vesicles. Intriguingly, we did not observe vesicles inside these clathrin cages, suggesting that clathrin protein molecules may spontaneously self-assemble into cages even when not involved in the endocytosis. It is important to note that the unexpected heptagon containing clathrin cage would be lost in averaging-based methods because it only has a single instance in the tomogram. Thus, these observations made in neurons demonstrated that IsoNet enables compensating for missing-wedge for structures that are highly heterogeneous, with limited copy numbers, and in the complex cellular environment.

## Discussion

Here we have developed a deep learning-based package, IsoNet, to overcome the limitation of missing-wedge problem and low SNR plaguing all current cryoET methods. IsoNet embodies several measures that prevent the neural network from “inventing” molecule features. First, the neural network was initialized with random numbers, and all the information comes from original tomograms without prior knowledge. Second, we introduced the dropout factor of 0.5 in the neural network so that with 50% of randomly picked neurons remaining, the network can still reproduce the result. Third, to further prevent over-fitting, the extracted subtomograms for training can be divided into random halves, and the resolution estimation is based on the gold-standard 3D FSC.

To demonstrate its robustness, we have applied IsoNet to process three representative types of cryoET data—pleomorphic virus HIV, cellular organelle axoneme with PFR, and neuronal synapse—representing three levels of length and complexity. IsoNet significantly improved structural interpretability in all these cases, allowing us to resolve novel structures of lattice defects in immature HIV capsid, dynein subunits, and scissors-stack-network architecture of the paraflagellar rod in eukaryotic flagella, and heptagon containing clathrin cage inside a neuronal synapse. In the resulting tomograms, the *in situ* protein features appear isotropic and have high quality that sometimes matches that obtained through subtomogram averaging. For amorphous structures in the tomograms, such as membranes, IsoNet allows the network to learn the feature representation from many other similar structures in the tomogram and recover the missing information. Thus, IsoNet expands the utility of cryoET by overcoming its inherent missing-wedge problem, enabling 3D visualization of structures that are either complex as those in cells (Figs. 4 and 5) or are rare as those tomograms of patients tissues^40^.

Philosophically speaking, no information can emerge from vacuum/nothing. So where does IsoNet recover the missing information from? The questions touch upon the fundamentals of deep learning and can be thought of as relating to non-locality of information in space. That is, by learning from information scattered around in original tomograms with recurring shapes of molecules, IsoNet sophistically eliminates distorted or missing information. The great advantage of the IsoNet approach is that similar features across different dimensions can be automatically discovered and “averaged” without human intervention. Such features could be related in translation and rotation manners in the three Cartesian dimensions, such as crystalline PFR subunits and axonemal microtubules and dyneins (Fig. 3); they could also be related through symmetries, such as those pentagons and hexagons of clathrin cages; ultimately, they could also be related biologically, such as the facts that proteins are made up of only 20 amino acids and nucleic acid of four bases, and both are geometrically constrained as a linear molecule. IsoNet learns their relationships in the same tomogram or across multiple tomograms and reconstructs these features automatically. In essence, therefore, IsoNet and sub-tomogram averaging compensate the missing-wedge problem through the same principle.

Regardless of the details of information recovery, the substantial improvement in map interpretability afforded by IsoNet now allows visualization of structures for functional interpretation without the need of tedious and time-consuming sub-tomogram averaging, which typically involves *a priori* feature identification and manual particle picking. Visualizing such structures in cellular tomograms by IsoNet would also improve localization and subsequent sub-tomogram averaging of hundreds of thousand copies of like-structures, leading to *in situ* atomic resolution structures of cellular complexes in their native cellular environment.

## Methods

### Software implementation

We implemented IsoNet in Python using Linux as the native operating system. Typical hardware setup includes one node with four Nvidia GeForce 1080Ti GPU cards of 11 gigabytes memory, which is common in a cryoET research laboratory. The package can be run from command line and relies on Keras that acts as interface for Tensorflow^15^, and the package can be downloaded from Github (https://github.com/Heng-Z/IsoNet). A detailed document is provided, accompanied by the IsoNet software. Tutorial dataset and video can be found in: https://github.com/Heng-Z/IsoNet_tutorial.

This package is standard-alone and does not rely on other software such as IMOD, while some common Python modules must be installed prior to running IsoNet. Such Python modules are easy to install with the “pip” command. For example, the IsoNet uses Python module “mrcfile” for the read and write tomogram or subtomogram, and “numpy” for the image processing such as rotation and Fourier transform. The U-net neural network is built by stacking multiple layers (Supplementary Fig. 3) provided in “tensorflow.keras.layers”. For example, three “Conv3D” layers are stacked together in each depth of the encoding path of the U-net.

The package can be launched through a single command entry, either “*isonet.py*” or “*isonet.py gui*”, for Linux command line operations or a graphical user interface (GUI) (Fig. 1b), respectively. Users can then access all the processing steps of the IsoNet procedure. The IsoNet procedure contains five steps, including three major steps: Extract, Refine, and Predict and two accessary steps: CTF deconvolve and mask generate. Each of these steps corresponds to one command of IsoNet in Linux terminal and will be described in the following sections.

### Dataset preparation

To use IsoNet, users should prepare a folder containing all tomograms. Binning the tomograms to a pixel size of about 10Å is recommended. Typically, a folder containing 1 to 5 tomograms is used as input. These input tomograms can either be reconstructed by SIRT or WBP algorithm. The tilt axis is the Y-axis, and recommended tilt range is -60 to 60°, while other tilt ranges might also work. The tilt series can be collected with any tilt schemes, continuous, bidirectional, or dose-symmetric.

IsoNet uses Self-defining Text Archive and Retrieval (STAR) file format to store information of tomograms and program parameters been used during Isonet processing. Thus, it inter-operates seamlessly with such leading cryoEM software packages as Relion^41^. Tomogram STAR file, default named as *tomograms.star*, is required to run IsoNet and this file can be prepared with IsoNet GUI, with a text editor, or with the following command:

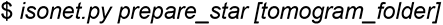

### Deconvolve CTF

For the tilt series imaged without Volta phase plate (VPP), the sinusoidal CTF suppresses or even inverts information at certain frequencies. To enhance the contrast of the tomograms and promote information retrieval, CTF deconvolution, similar to what is implemented in Warp software, is applied to tomograms in this step.

IsoNet uses Wiener-like filter^16^ for CTF deconvolution, with spectral signal-to-noise ratio (SSNR) set empirically:

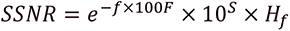

Where f denotes the spatial frequency, H a high-pass filter, F the custom fall-off parameter, and S denotes the custom strength parameter. Because the SSNR of the Weiner-like filter is determined empirically, users can tune the SSNR fall-off or deconvolve strength parameters to enhance contrast of the tomograms. This step can be performed by “CTF deconvolve” function in GUI or with the following command:

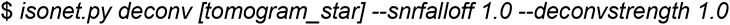

### Generate mask

Subtomograms for training would be better to contain rich information than empty areas with only ice, air, or carbon. In this optional mask generation step, IsoNet uses statistical methods to detect empty regions from which subtomograms will not be extracted. Two different types of masks can be applied: density mask that excludes areas with low cryoET density and standard deviation mask that excludes areas with low standard deviation.

The density mask will first suppress the noise with a Gaussian filter and then apply a sliding window maximum filter to the contrast-inverted tomogram. The areas with relatively smaller density values in the filtered tomogram will be deemed as empty space and excluded in the mask. The parameter “*density_percentage*” defines the percentage of the area kept in the tomogram by the density mask.

The standard deviation mask is achieved by calculating the standard deviation of voxels in a small cubic volume centered at each evaluating voxel. The voxels having relatively lower standard deviation will be excluded. The parameter “*std_percentage*” defines the percentage of voxels kept by the standard deviation mask.

IsoNet uses the intersection of these two masks. The parameters for mask generation can be tuned to cover the region of interest but exclude empty areas. In addition to these two types of masks, IsoNet allows excluding top and bottom parts of tomograms which usually are empty areas by the “*z_crop*” parameter.

Usually, the default parameters will provide a good mask to exclude empty areas; users can also tune the parameters using GUI by “Generate mask” function or the following command, for example:

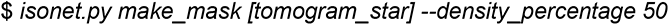

### Extract subtomograms

In each tomogram, specified number of seeds are randomly generated within the whole tomogram or the region of interest defined by the mask. Then, cubic volumes centered at the generated seeds are boxed out and saved as subtomograms. The extracted subtomograms should be large enough to cover typical features in tomograms, such as a patch of membrane or vesicle. However, due to the GPU memory limitation, this size cannot be arbitrarily large. We usually extract 300 subtomograms of 96^3^ voxels in total. After extraction, the contrast of those subtomograms is inverted. Then, tomograms are normalized by percentile to ensure 80% of the voxel values fit into the range between zero and one. The subtomograms can be randomly split into two halves and used for performing missing-wedge correction independently to eliminate overfitting and calculate gold-standard FSC.

The information of extracted subtomograms is stored in another STAR file, default named as *subtomo.star*. Subtomogram extraction can be performed by either IsoNet GUI or the following command:

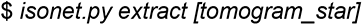

### Refine

This process iteratively trains neural networks to fill the missing-wedge information using the same tomograms whose missing-wedge artifacts were added to other directions. The denoising module can also be enabled in this step, making the network capable of reducing noise and recovering the missing-wedge. After refinement, the resulting subtomograms and neural network model in each iteration are saved. The network models with a suffix of “.h5” can be used for the prediction step.

Four steps, including *training dataset generation*, *adding noise*, *network training*, and *subtomograms prediction*, will be performed during each iteration. These steps will be described in the following sections. The missing-wedge restored subtomograms by *subtomograms prediction* in every iteration will be used for *training dataset generation* in the next iteration. Usually, 10-15 iterations in the refine step are sufficient to obtain a well-trained network for the missing-wedge correction, whereas more iterations can be performed for refinement with denoising.

The refine step can be performed from the GUI or with the following command, for example:

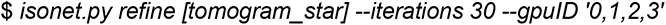

Users can also continue training from the previous iterations using “*continue_from*” or from previously trained models using “*pretrained_model*” parameter.

### Refine step 1: training dataset generation

To generate paired datasets for neural network training, IsoNet rotates the extracted subtomograms to different orientations. Twenty rotated copies can be obtained for each extracted subtomogram as follows (Supplementary Fig. 2). First, each subtomogram is a cube with six faces. Each face can be rotated with an out-of-plane angle to face toward the positive direction of the Z-axis. Second, each out-of-plane rotation can be followed by four in-plane rotations, making 24 possible rotations. However, among the 24 rotations, four of them result in subtomograms with the same missing-wedge direction as the original subtomograms. Thus, these four rotations are excluded, resulting in 20 orientations for each subtomograms. This rotation process enlarges the original dataset by 20 times for training, making it possible to achieve a good performance of missing-wedge correction even with a small dataset, *e.g.,* a single tomogram.

After the rotation, the IsoNet program then applies missing-wedge filter to the rotated subtomograms. The missing-wedge filter volume has the same size as that of the subtomograms. In the missing-wedge filter volume, voxel value is zero inside the missing-wedge region and one in the rest of the volume. Then, the Fourier transforms of the rotated subtomograms are multiplied by the missing-wedge filter volume and then transformed back to the real space, generating missing-wedge filtered subtomograms.

To avoid the incomplete information along the edge of the subtomograms when applying missing-wedge filter, both rotated subtomograms and missing-wedge filtered rotated subtomograms are trimmed into smaller volumes (often 64^3^ voxels), generating “target” and “input” for the network training, respectively. These generated data pairs are used to train neural network that maps the “input” to “target”.

### Refine step 2: adding noise

This optional step allows performing missing-wedge correction and denoising simultaneously using IsoNet. IsoNet uses a noisier-input strategy^20,21^ that learns to map “input” with additional noise to the “target”.

IsoNet simulates the noise pattern in reconstructed tomograms with the assumption that in every acquired projection, the noise is an additive Gaussian noise and independent among all images acquired in a tilt series. During the adding noise step, a set of 3D noise volumes are constructed by back-projecting a series of 2D Gaussian noise images to reflect the effect of the back-projection algorithm on noise formation.

The denoise level is defined as the ratio of the standard deviation between the subtomograms and the added noise. The noise volumes are scaled to match the denoise level before being added to the “input” subtomograms. Thus, the lower denoise value means less noise is added to individual subtomograms.

Because the added noise may further corrupt the 3D subtomograms, making the network hard to train, it is recommended to start the first several iterations of refinement without denoising. After the refinement results are nearly converged, the noise volume can then be added to the “input” subtomograms in the following iterations. A typical routine is to train ten iterations without denoising and then increase the denoise level by 0.05 for every five iterations. This step-wised noise addition can be performed automatically in the refine step of the IsoNet software.

### Refine step 3: network training

Neural network used in IsoNet is based on U-net, which is well recognized in biomedical image restoration and segmentation^19^. The main building blocks of the U-net are 3D convolution layers with non-linear activation functions called Rectified Linear Units (Relu), which are applied per voxel. Those convolution layers have kernel sizes of 3×3×3. Three 3D convolution layers are stacked together to form a convolution block in our network, which can extract complicated features.

By stacking the convolutional blocks, the U-net is built based on encoder-decoder architecture (Supplementary Fig. 3). The encoder path is a set of convolution blocks and strided convolution layers that compress 3D volumes. Strided convolution layers reduce the spatial size of the input of this layer by 2×2×2, allowing the network to learn more abstract information. A convolution block followed by a strided convolution layer makes one encoder block in the contracting path. Total three encoder blocks form the entire encoding path. The number of convolution kernels for each convolution layer doubles after each encoder block. After the encoder path, the 3D volumes are processed with a convolution block and enter the decoder path of the network. The decoder path is symmetrical to the encoder but uses transpose convolution layers, opposite to strided convolution layers, to enlarge the dimension of features.

Although the down-sampling of the 3D volumes captures the essence of the features, high-resolution information is lost by stride convolution operations. In particular, the skip connections that concatenate the feature layers of the same dimension in two paths are implemented to preserve the high-resolution information. Dropout strategy that randomly set 50% of neurons’ activation to 0 in the convolution layers is used to prevent overfitting during the training.

This network uses the mean absolute error between the output of the network and the target subtomograms as loss function. The loss function is minimized by employing Adam optimizer^42^ with an initial learning rate of 0.0004. The neural network training is performed on GPU and consists of several epochs or cycles (typically ten epochs). Each epoch will traverse through the paired dataset. The data pairs are grouped into batches (which generally have a size of 8 or 16) to feed into each epoch. After the training, the trained neural networks are saved for the next iteration of the refine step.

### Refine step 4: Subtomogram prediction

After each iteration of refinement, the network is applied to the original subtomograms, generated predicted subtomograms. Then IsoNet generates an inverse missing-wedge filter volume with value one inside the missing-wedge region but zero in the rest of the 3D volume. The predicted subtomograms are then transformed to Fourier space and multiplied with the inverse missing-wedge filter volume to extract the added information inside the missing-wedge regions. Then, these filtered volumes are added to the original subtomograms, generating the missing-wedge restored subtomograms for subsequent iterations of refinement.

### Predict

After the refine step, the trained network is saved in a model file. It will be used to correct the missing-wedge for the original tomogram or other similar tomograms. For most tomograms, the full-size 3D images can hardly fit into the memory of a regular GPU. Thus, the IsoNet program splits the entire tomogram into smaller 3D chunks to apply the network on them separately. Then output 3D chunks are montaged to produce the final output. To avoid the line artifact between adjacent chunks caused by the loss of information on the edges of subtomograms. We implemented a seamless reconstruction method called overlap-tile strategy, which predicts the overlapping chunks to avoid the edge effect. The “crop_size” parameter defines the size of the cubic chunks. This predicting step can be performed with IsoNet GUI or with the following command, for example:

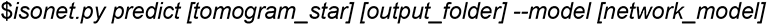

### Benchmarking with simulated data

We performed IsoNet reconstruction on simulated subtomograms using the public available atomic models: apoferritin model^22^ (PDB: 6Z6U) and ribosome model^23^ (PDB: 5T2C). For both tests, density maps were simulated from the atomic models using “*molmap*” function in ChimeraX^24^ and filtered to 8Å resolution (Fig. 1d, e). Those simulated maps with 2.67 Å/pix pixel size were then rotated in 10 random directions and imposed with missing-wedge filter in Fourier space, resulting in simulated subtomograms with missing-wedge artifacts (leftmost columns in both Fig. 1d, e).

For the simulated Apoferritin maps, we created a subtomogram STAR file with the “*isonet.py prepare_subtomo_star*” command. With this subtomogram STAR file as input, we performed an IsoNet refine step for ten iterations without denoising. For benchmarking with the simulated ribosome maps, we extracted eight smaller subtomograms (96^3^ voxels) from each ribosome map due to the larger dimension of the ribosome map. The subtomogram STAR file generated in the extract step was used for subsequent refine step. After ten iterations, trained network was obtained and was then used to produce missing-wedge corrected maps of ribosome using “isonet.py predict” command.

### Processing tomograms of HIV virus

For pleomorphic viruses, we downloaded an HIV dataset from public repository EMPIAR-10164^26^. Three tilt series,TS_01, TS_43 and TS_45, were used for testing. The movie stacks were drift corrected with MotionCorr^43^ and reconstructed with IMOD^11^ using WBP algorithm. The defocus value of each image was determined by CTFFIND4^44^. Eight-time binned tomograms with 10.8Å pixel size were used for further processing. For the CTF deconvolution of the tomograms, SSNR fall-off and the deconvolve strength parameters were set to 0.7 and 1, respectively. Then, we created one mask for each tomogram using “isonet.py make_mask” command. Total 300 subtomograms with 96^3^ voxels were randomly extracted from the three tomograms and then split into random halves. For each half of the subtomograms, we performed refine step for 35 iterations independently, generating two trained neural networks. In the predict step of IsoNet, tomogram TS_01 was used to generate two missing-wedge corrected tomograms by the two independently trained networks. These two tomograms were then averaged to create a final map.

These two missing-wedge corrected tomograms enabled calculating gold-standard FSC. Instead of calculating a global FSC, we performed 3D FSC calculation for all the directions^17^ to measure the resolution anisotropy of the missing-wedge corrected tomogram. Because the 3D FSC calculation works for cubic volumes while the size of the tomogram is non-cubical, we cropped the generated tomograms into cubic subtomograms for the 3D FSC calculation. As for the HIV dataset, the 3D FSC was calculated for four 200^3^ volumes cropped from both missing-wedge corrected HIV tomograms. The resulting four 3D FSC were then averaged to produce the final 3D FSC, the orthogonal sections of which are shown in Figure 2b.

### Processing tomograms of the Eukaryotic flagella

For cellular organelles, we chose the demembraned flagella of *Trypanosoma. Brucei.* The datasets described here were obtained in our previous studies^32,45^. Tilt series were recorded with SerialEM^46^ by tilting the specimen stage from −60° to +60° with 2° increments. The cumulative electron dosage was limited to 100 to 110 e^−^/Å^2^ per tilt series. The movie stacks were drift corrected with MotionCorr^43^ and reconstructed with IMOD using SIRT algorithm. The tomograms were binned by four, resulting in a pixel size of 10.21 Å/pixel.

Three tomograms were chosen for missing-wedge correction. These tilt series were acquired with VPP so that we did not perform the CTF deconvolution. We generated one mask for each tomogram using “isonet.py make_mask” command. Then, we extracted a total of 360 cubic subtomograms with 128^3^ voxels from three tomograms. Using these subtomograms, we trained a network by running the refine step for 35 iterations with default denoise levels, which were automatically changed from 0 to 0.2. Trained network produced in the refine step was then used to run the predict step of IsoNet to obtain a final missing-wedge corrected tomogram, which is shown in Figure 3.

### Processing tomograms of hippocampal neurons

Tomograms of hippocampal neurons were obtained in our previous study^7^. The two tomograms used in this study were collected on a Titan Krios microscope equipped with K2 summit in counting mode. The energy filter (Gatan image filter) slit was set at 20 eV. The Titan Krios was operated at an acceleration voltage of 300 KV. Tilt series were acquired using SerialEM^46^ with tilt scheme: from +48° to -60° and from +50° to +66° at an interval of 2°. The total accumulated dose was ∼150 e-/Å^2^. The pixel size of the tomograms is 4.35 Å/pixel. Each recorded movie stack was drift-corrected and averaged to produce a corresponding micrograph using MotionCorr^43^. The tilt series were aligned using IMOD^11^. One tilt series of the tomogram shown in Figure 4 was imaged with VPP, while the other shown in Figure 5 was acquired without VPP. When VPP was used, the defocus value was maintained at -1 μm; otherwise, it was kept at -4 μm.

For the tilt series recorded with VPP, the aligned tilt series were reconstructed using NovaCTF^47^, generating a tomogram reconstructed with WBP. The tomogram was binned by four, and 300 subtomograms (96^3^ voxels) were extracted from that tomogram. Those subtomograms were then used 35 iterations of refinement. The trained network produced in refine step was used for missing-wedge correction for the entire tomogram (Fig. 4).

For the tilt series recorded without VPP, the defocus value of each image was determined by CTFFIND4^44^, and CTF phase flipped tomogram was obtained by NovaCTF^47^. This tomogram (Fig. 5) was reconstructed with SIRT-like filter, with CTF phase flipping performed on the individual tilt images. The tomogram was binned by four for missing-wedge correction with IsoNet. Then, 200 subtomograms (96^3^ voxels) were extracted from the tomogram in the extract step of IsoNet. A trained network was obtained with the refine step of IsoNet for 35 iterations. The trained network was then used for the predict step of IsoNet, producing a missing-wedge corrected tomogram (Fig. 5).

### 3D visualization

IMOD^48^ was used to visualize the 2D tomographic slices. UCSF ChimeraX^24^ was used to visualize the resulting IsoNet generated tomograms in their three dimensions. Segmentation of densities maps and surface rendering were performed by the volume tracer and color zone in UCSF ChimeraX.

## Supplementary information

Supplementary Information is linked to the online version of the paper.

## Acknowledgments

We thank Chong-Li Tian, Yi-Nan Xia and Zhuo Li for testing the program, and Jiayan Zhang, Simon Imhof, Ivo Atanasov, Wong Hui and Kent Hill for the Eukaryotic flagellum data. This work was supported in part by grants from the National Key R&D Program of China (2017YFA0505300), the National Natural Science Foundation of China (31630030, 31761163006, and 31621002), the Strategic Priority Research Program of the Chinese Academy of Sciences (XDB32030200). Research in the Zhou group is supported in part by the U.S. National Institutes of Health (GM071940). We acknowledge the use of resources at the Center for Integrative Imaging of Hefei National Laboratory for Physical Sciences at the Microscale, and those at the Electron Imaging Center for Nanomachines of UCLA supported by U.S. NIH (S10RR23057 and S10OD018111) and U.S. NSF (DMR-1548924 and DBI-133813).

## Author contributions

Y.-T.L. conceptualized the method, wrote code, processed data, made illustrations and documentation, and wrote the paper; H.Z. wrote code, processed the HIV and neuronal synapse data, made documentation and assisted in illustrations, and wrote the paper; H. W. participated in coding, processed the axoneme with PFR data, made documentation and assisted in illustrations; C.-L.T. assisted in testing the method with the neuronal synapse data and in making illustration based on the test results; G.-Q. B. and Z.H.Z. oversaw the project, interpreted the results, and wrote the paper. All the authors edited and approved the manuscript.

## Author information

The authors declare no competing financial interests. Correspondence and requests for materials should be addressed to G.-Q.B. or Z.H.Z.

## Data and materials availability

All data reported in the main text or the supplementary materials are available upon request. The software has been deposited to GitHub page at https://github.com/Heng-Z/IsoNet.

**Supplementary Fig. 1.**
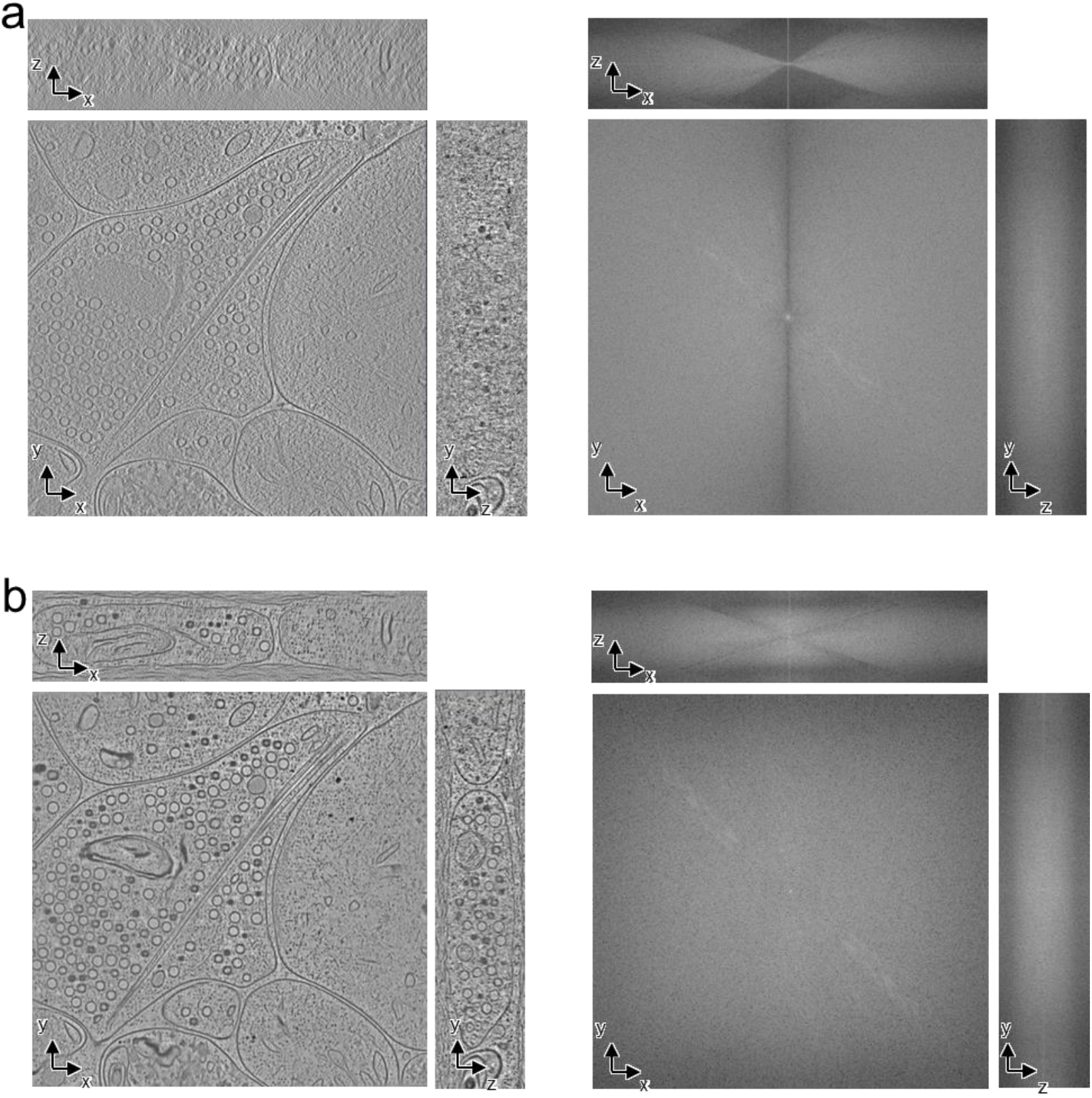
**a,** Orthogonal views of the WBP reconstructed tomograms and their corresponding Fourier transforms. **b,** Orthogonal views of the IsoNet reconstructed tomograms and their corresponding Fourier transforms.

**Supplementary Fig. 2.**
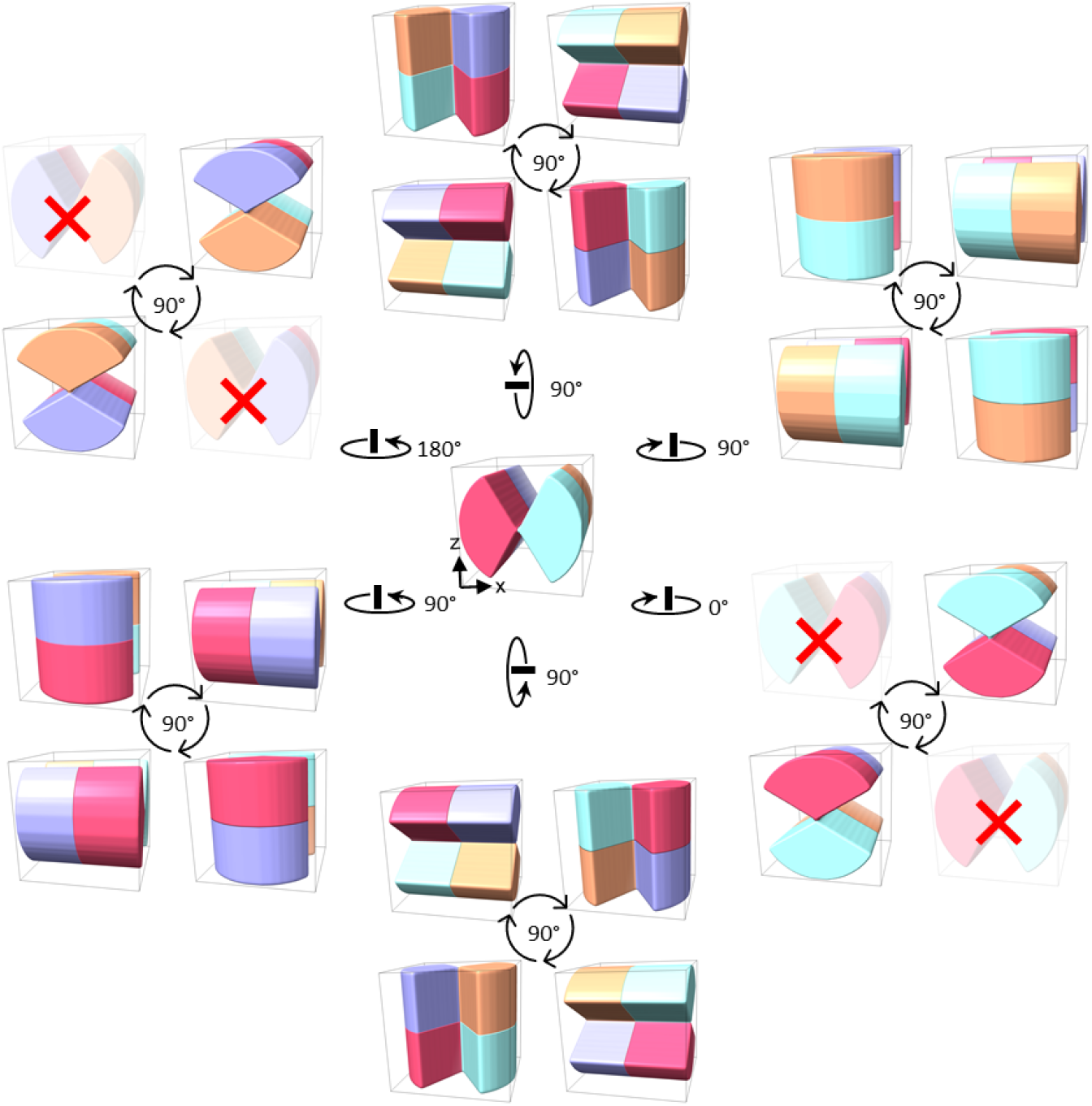
Rotation schemes. Twenty rotated copies are obtained for each extracted subtomograms demonstrated in the center. First, each subtomogram has six faces. Each face can be rotated with an out-of-plane angle to face toward six positive directions of the Y-axis. Second, each out-of-plane rotation is followed by four in-plane rotations, making a total of 24 possible rotations. However, among the 24 rotations copies, four have the same Z-axis missing-wedge as the original subtomograms. Thus, these four rotations are excluded (red cross).

**Supplementary Fig. 3.**
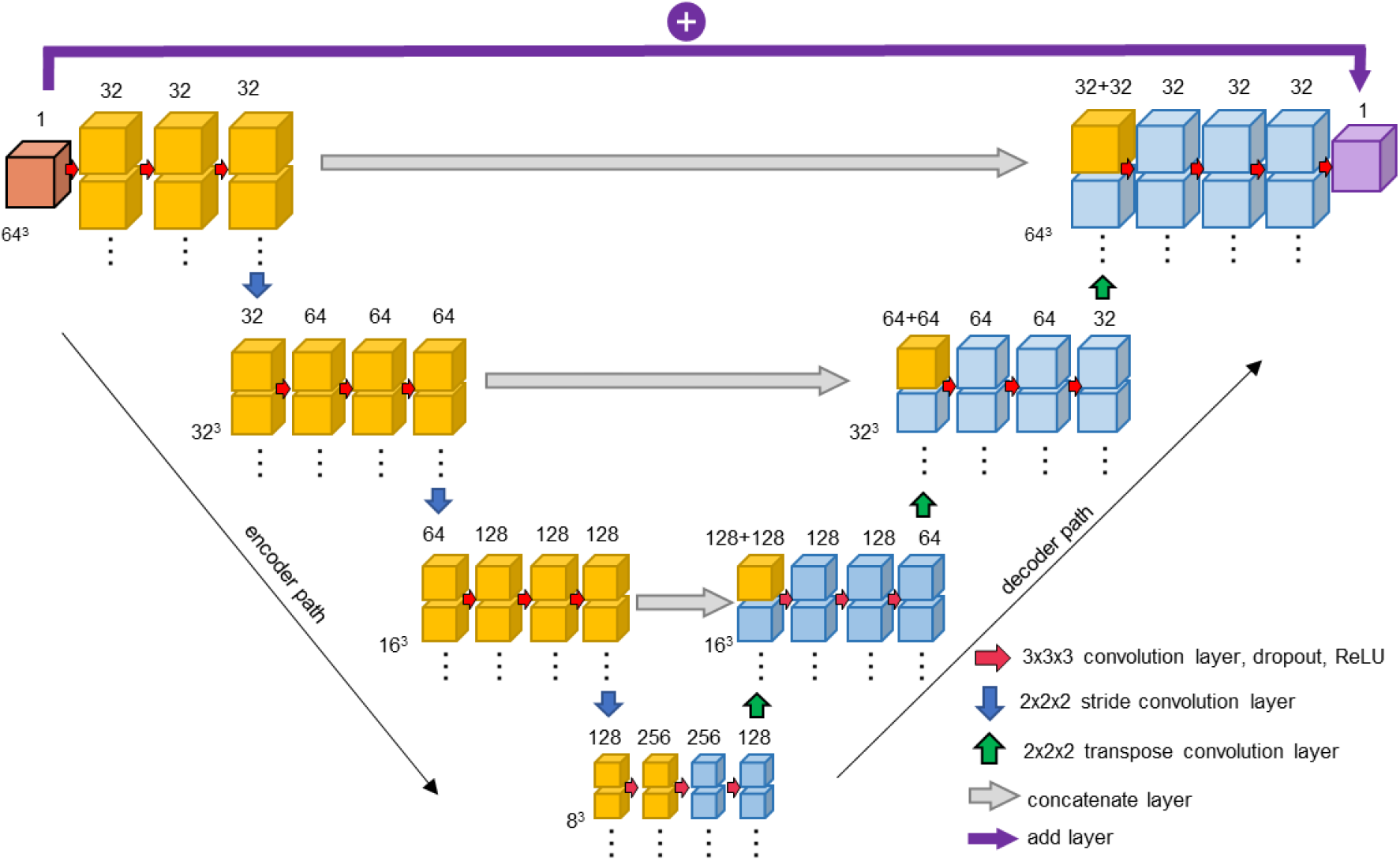
The architecture of neural network based on U-net. The values at the bottom left of boxes show sizes of 3D feature maps or subtomograms, while the values on top of the boxes are their numbers.

**Supplementary Fig. 4.**
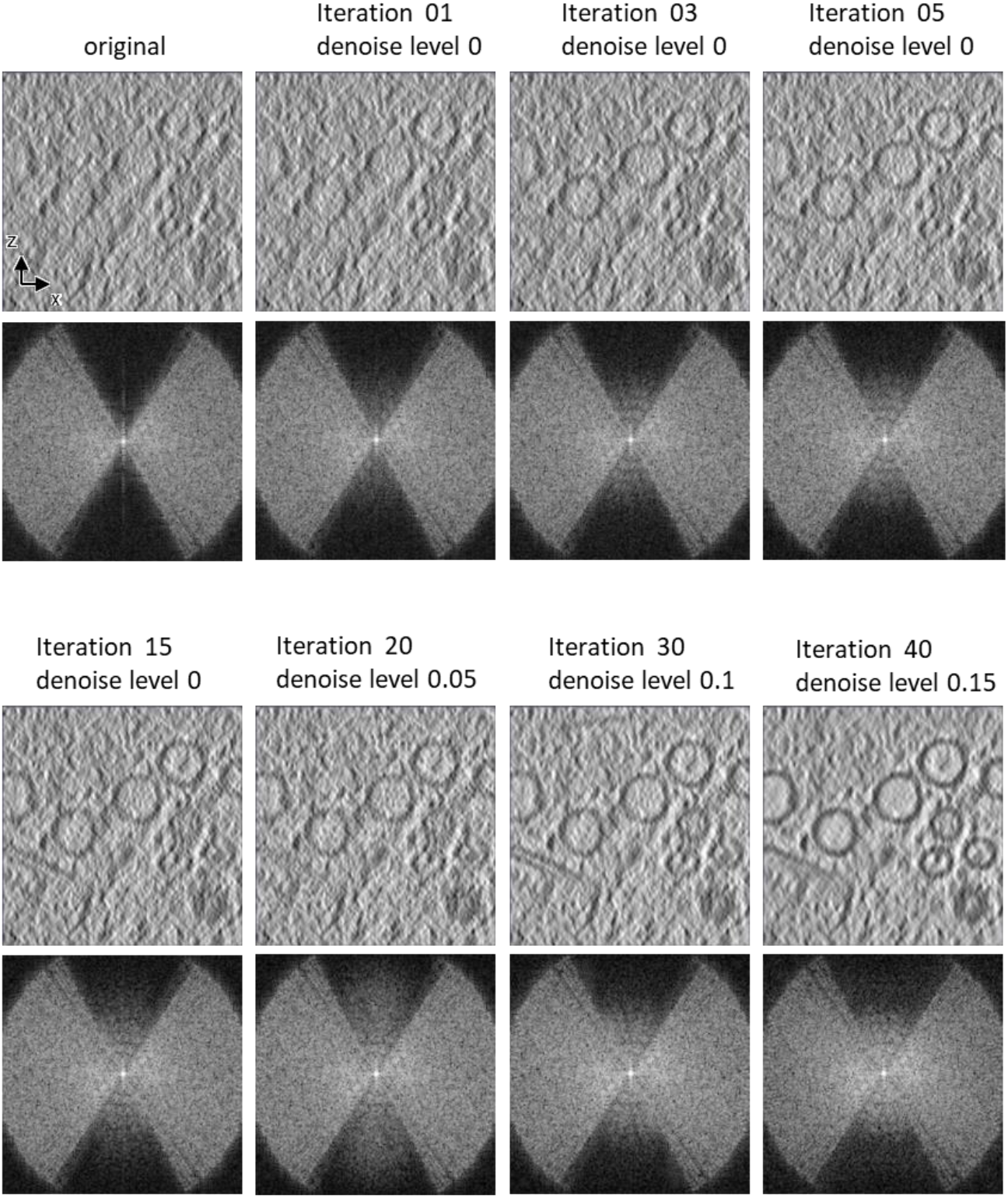
Iteratively filling the missing-wedge region. XZ slice views of subtomograms and corresponding power spectrum at different iterations in refine step.

**Supplementary Fig. 5.**
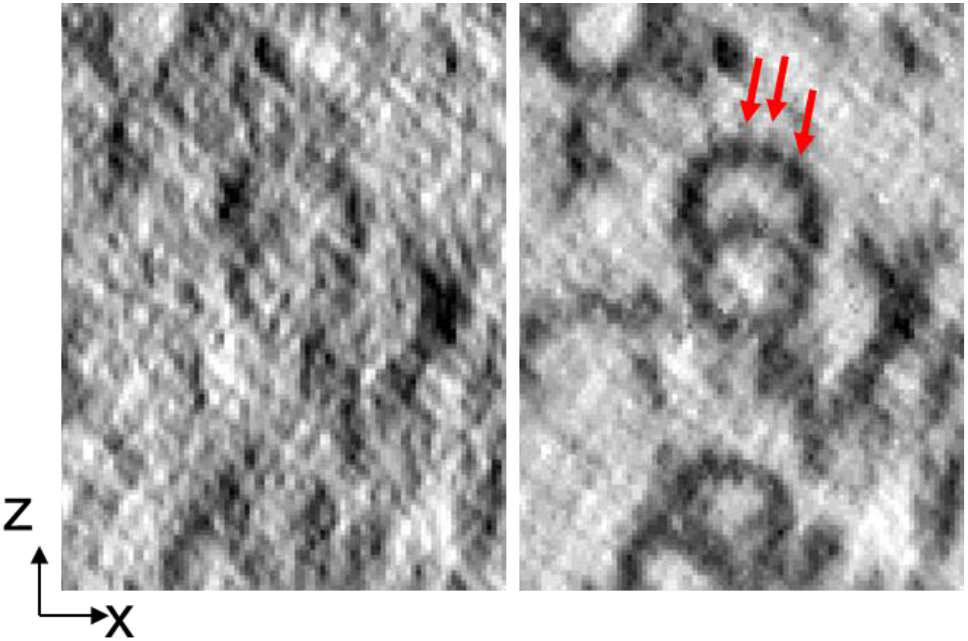
XZ slices of microtubule doublets in axoneme. Left: XZ slices of the SIRT reconstructions. Right: the slices of the corresponding tomogram generated by IsoNet. Red arrows indicate microtubule subunits.

**Supplementary Fig. 6.**
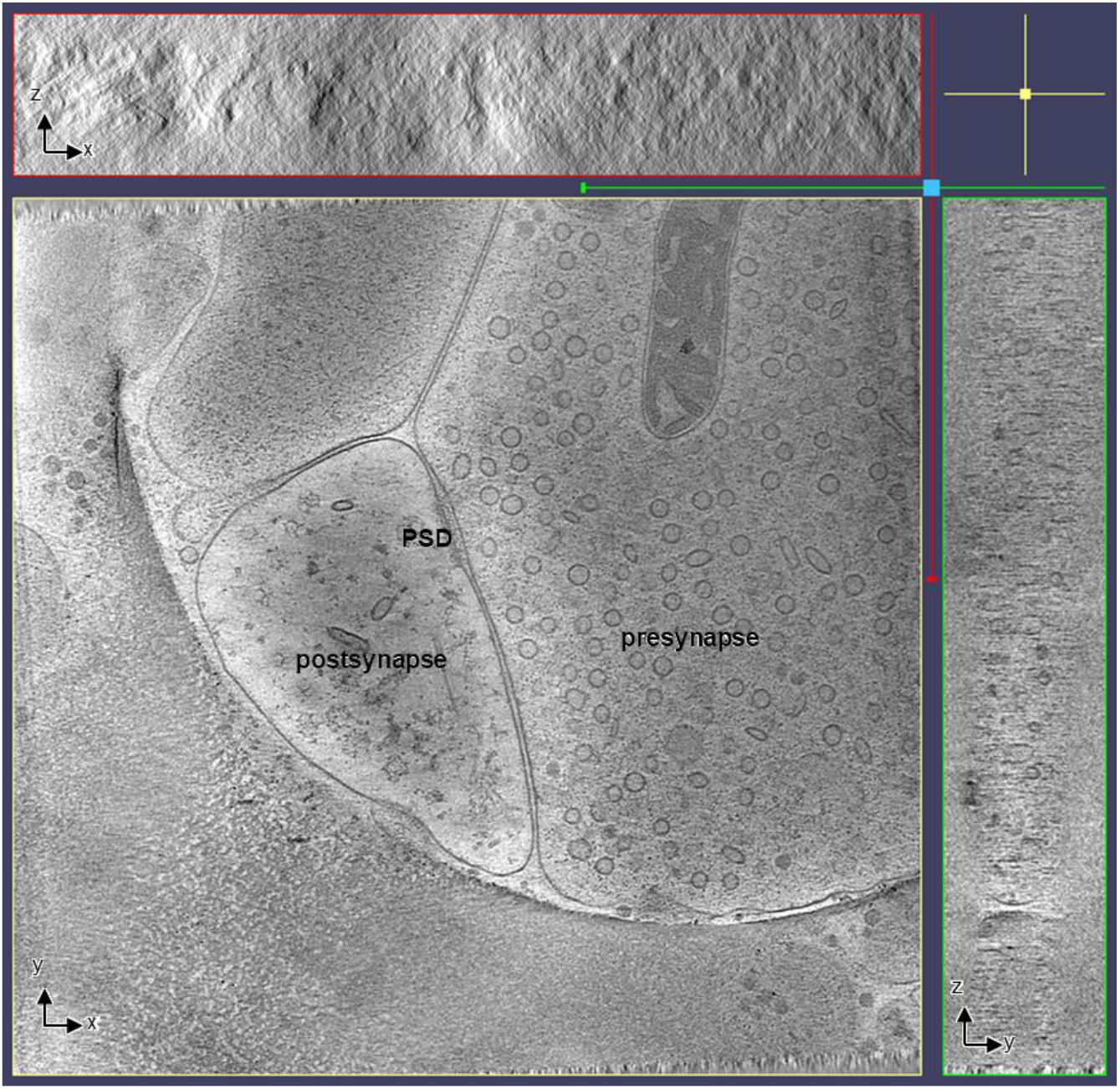
Orthogonal views of the tomogram containing clathrin cages reconstructed with the SIRT algorithm in IMOD. PSD: postsynaptic density.

**Supplementary Fig. 7.**
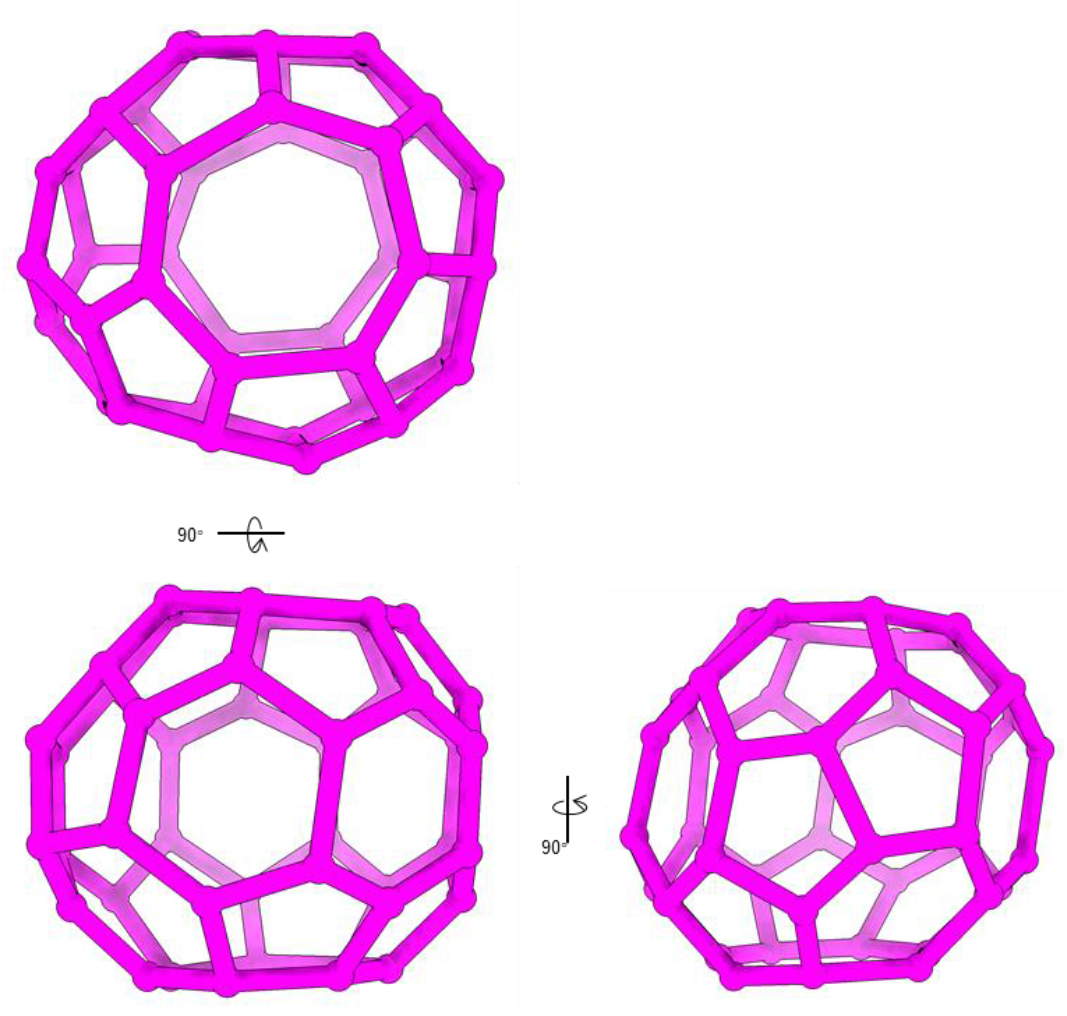
3D views for the shape of the heptagon containing clathrin cage.

**Supplementary Video 1.**
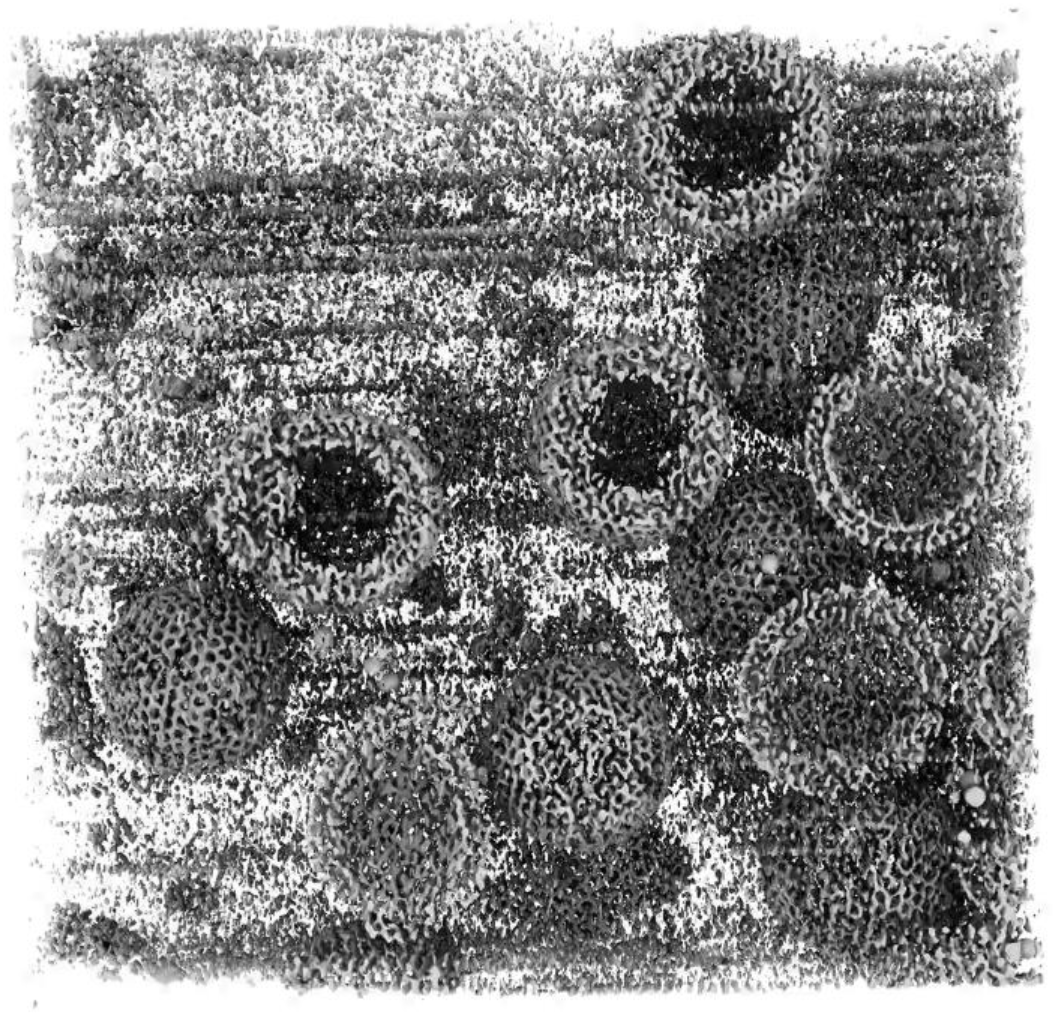
3D tomogram of HIV particles. This video shows 3D structures in an IsoNet generated tomogram. The tomogram density is sliced through three orthogonal directions.

**Supplementary Video 2.**
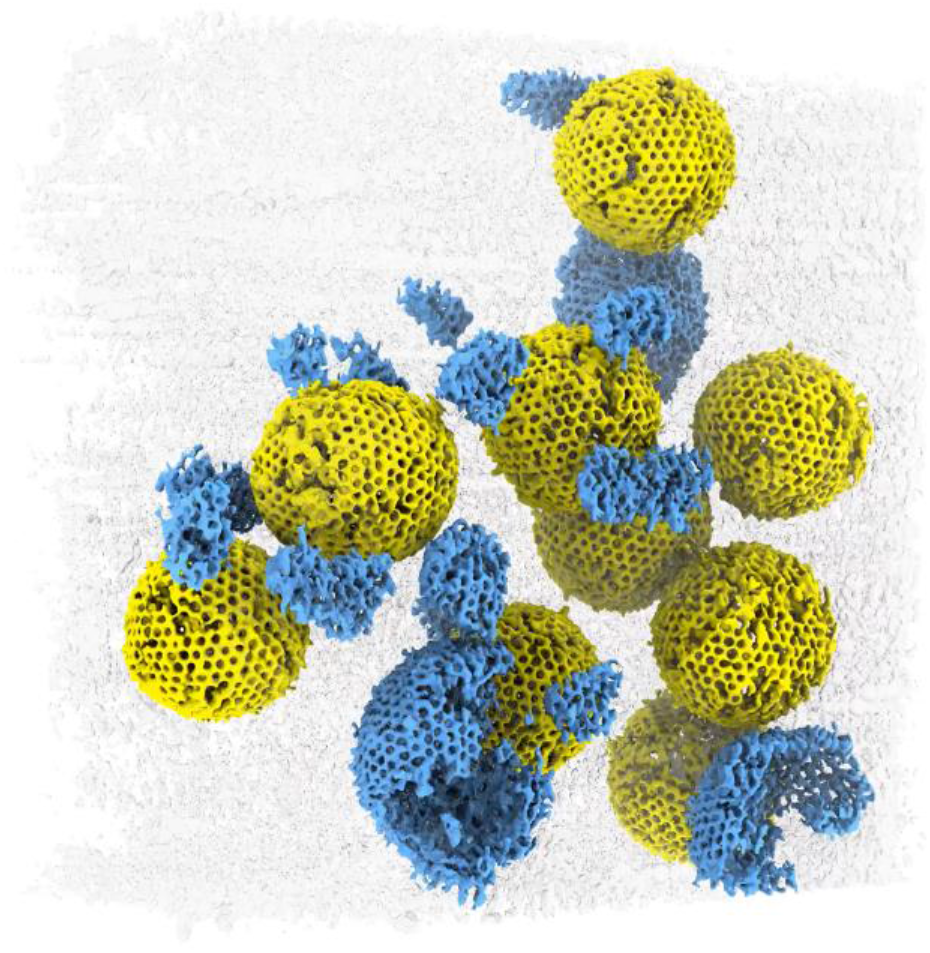
3D rendering of HIV particles. The HIV particles are rendered in yellow (fully embedded in ice) and blue (at air-water interface). The rest cryoEM density is shown in transparent gray.

**Supplementary Video 3.**
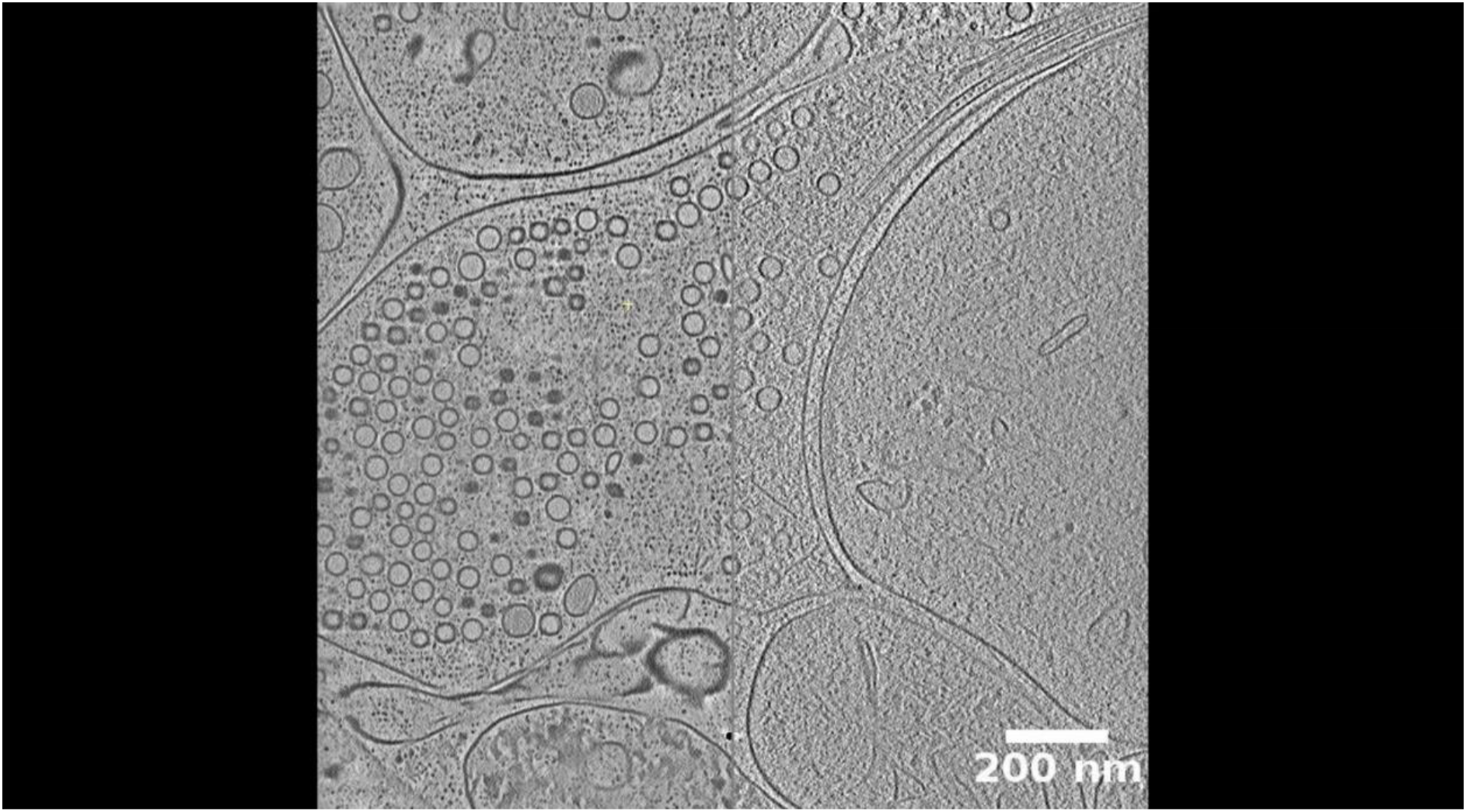
2D slices of a tomogram of a neuronal synapse. Left: IsoNet generated tomogram. Right: The original tomogram reconstructed with WBP.

**Supplementary Video 4.**
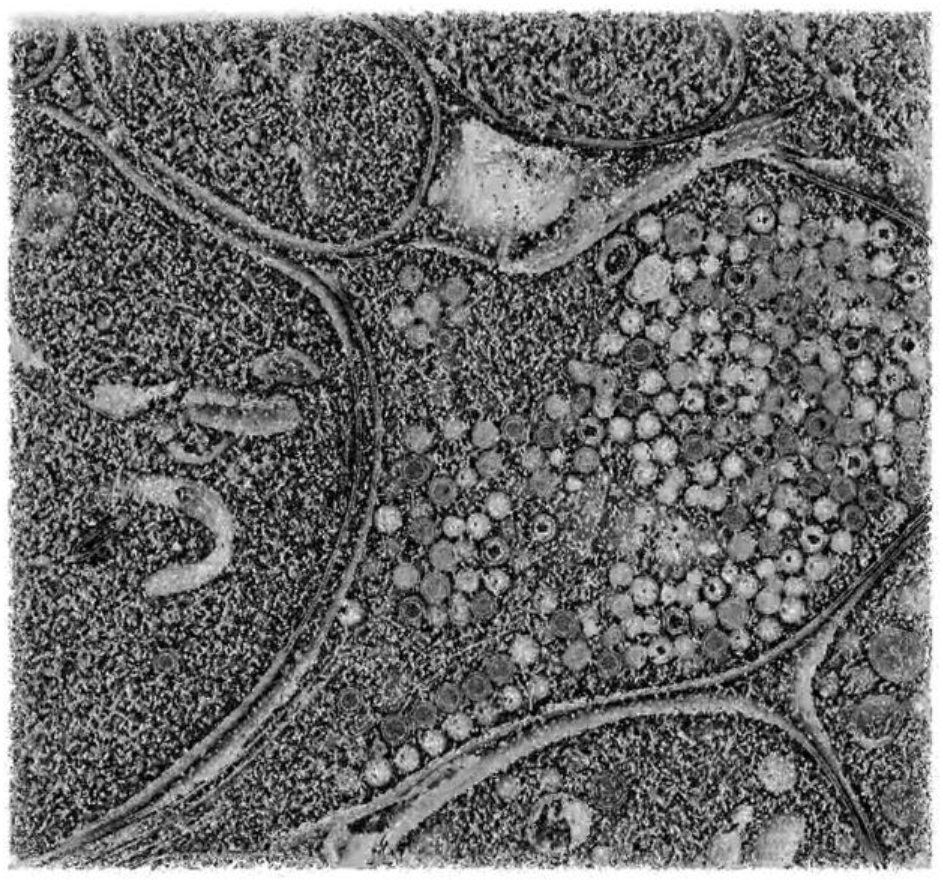
3D tomogram of a neuronal synapse. This video shows 3D structures in an IsoNet generated tomogram. The tomogram density is sliced through three orthogonal directions.

**Supplementary Video 5.**
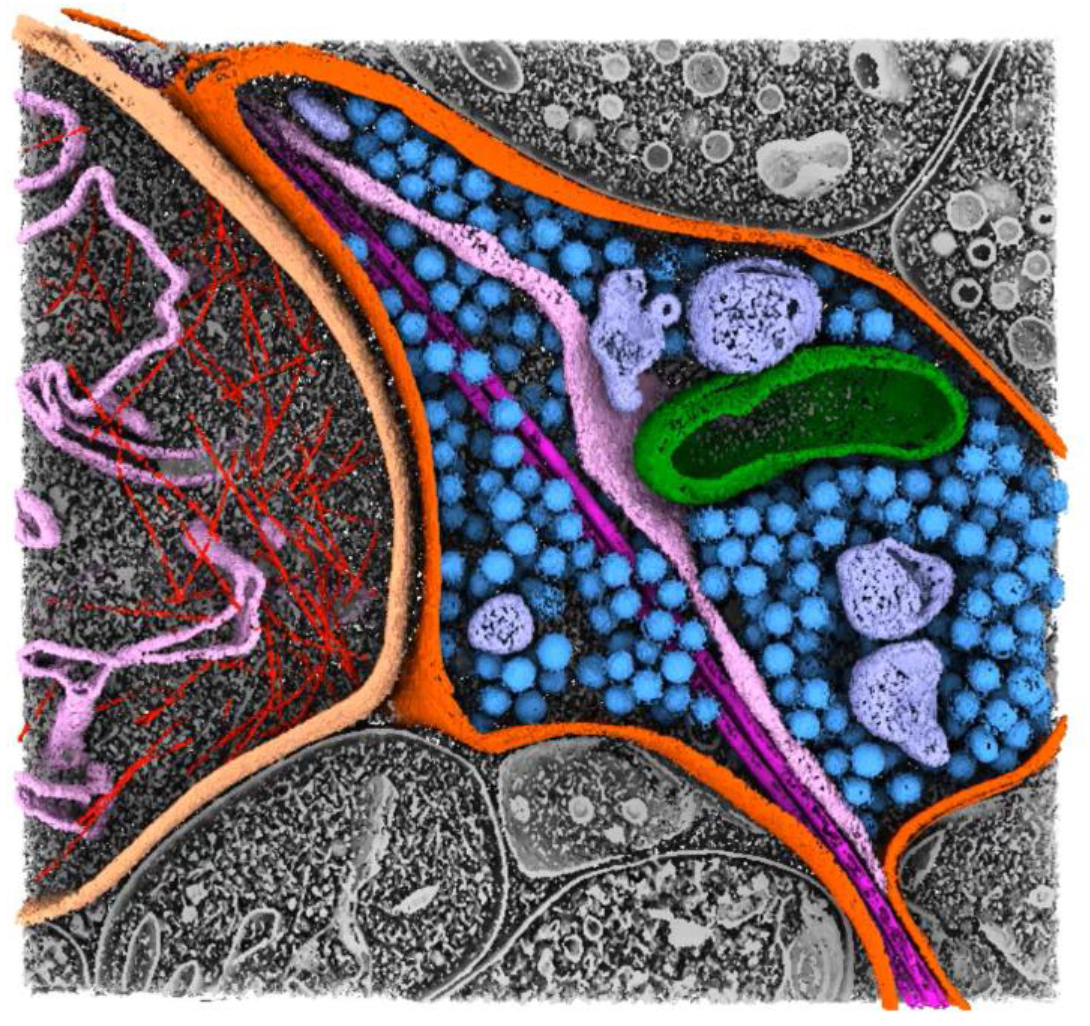
3D rendering of the neuronal synapse.

**Supplementary Video 6.**
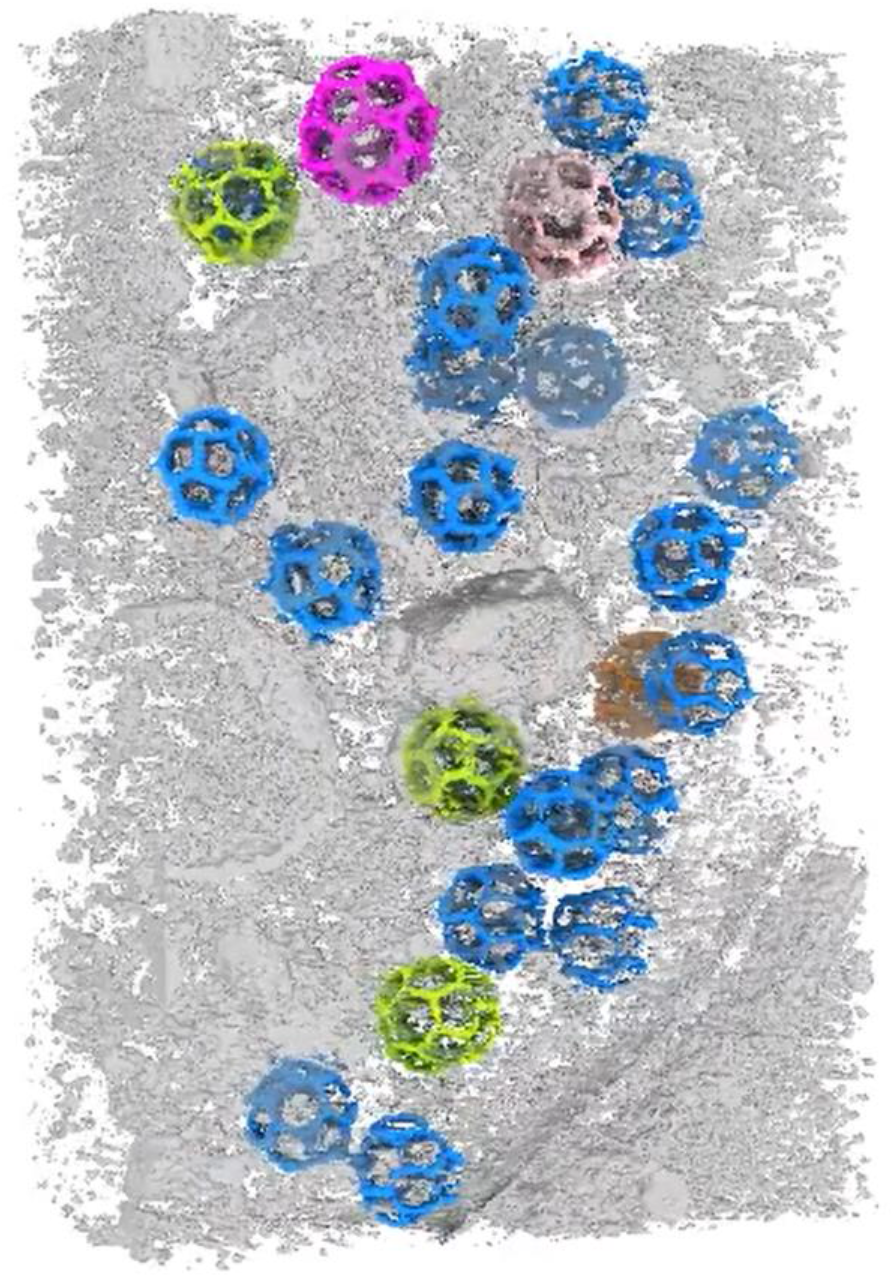
3D tomogram clathrin cages in a neuronal synapse. This video shows 3D structures of an IsoNet generated tomogram. The tomogram density is sliced through three orthogonal directions. Then, 3D rendering of clathrin cages is shown and rotated.

